# HV–MultiScaleGNN: A Multi-Scale Graph Neural Network Framework for Host–Virus Protein–Protein Interactions Prediction

**DOI:** 10.64898/2026.11.08.765214

**Authors:** Sarangthem Dinamani Singh, Gori Sankar Borah, Selvaraman Nagamani

## Abstract

Accurate identification of HV–PPIs is essential for understanding viral infection mechanisms and for developing antiviral therapeutic discovery. However, the experimental identification process is often time-consuming and expensive, while many computational approaches depend on incomplete protein sequences, structural and interaction information. Here, we present HV–MultiScaleGNN, a topology-driven multi-scale GNN framework for HV–PPIs prediction that operates exclusively on the topology of a heterogeneous interaction network. The framework formulates HV–PPIs prediction as a heterogeneous link prediction problem by integrating the human interactome with experimentally validated HV–PPIs. HV– MultiScaleGNN systematically investigates three complementary graph representation strategies, viz. Local view, Global view and an integrated Local+Global view. The Local view employs one-hop relation-aware graph attention to model fine-grained neighbourhood interactions. The Global view captures higher-order structural dependencies through propagation-based neighbourhood aggregation. The integrated representation combines both views to jointly encode local connectivity and global topological context within a unified embedding space, enabling relation-specific link prediction without relying on sequence- or structure-derived features. Experiments on large-scale HV–PPIs datasets evaluate all three strategies using discrimination, calibration, and ranking metrics. The Local view achieved the best discriminative performance (AUROC = 0.9444, AUPRC = 0.9422), whereas the Global view exhibited the lowest ECE (0.0379). The integrated Local+Global view provided consistently strong predictive performance (AUROC = 0.9378, AUPRC = 0.9356). Comparative evaluation against representative graph representation learning methods showed competitive predictive performance. Furthermore, interfacial interactions were identified using protein-protein docking and atomistic-level understanding was explored. Moreover, GO and KEGG enrichment provided additional functional insights into the selected host proteins. The proposed framework enables systematic evaluation of complementary graph representation strategies for heterogeneous biological link prediction and helps experimental researchers to delineate novel HV–PPIs.

## 1. Introduction

Viruses rank among the most abundant and genetically diverse biological entities and are responsible for a wide spectrum of infectious diseases affecting humans, animals, and plants. Their survival and replication depend on the ability to hijack host cellular machinery through highly coordinated molecular interactions [1]. Among these interactions, host–virus protein–protein interactions (HV–PPIs) play central roles throughout the viral life cycle, including viral entry, genome replication, intracellular trafficking, immune evasion, and virion assembly [2, 3]. Because viral proteins selectively interact with host proteins to manipulate cellular pathways and establish a favorable environment for infection [4]. Systematic identification of HV–PPIs is essential for understanding virus–host interactions, elucidating disease mechanisms, and accelerating antiviral drug discovery and therapeutic development [5].

Despite substantial advances in experimental biology, comprehensive identification of HV–PPIs remains a challenging task. High-throughput experimental techniques, including yeast two-hybrid (Y2H) [6], mass spectrometry (MS) [7], co-immunoprecipitation (co-IP) [8], fluorescence resonance energy transfer (FRET) [9], tandem affinity purification (TAP) [10], protein fragment complementation assays (PCAs) [11], and bimolecular fluorescence complementation (BiFC) [12] have considerably expanded the discovery of protein–protein interactions (PPIs). Nevertheless, these approaches remain labor-intensive, time-consuming, technically demanding, and susceptible to both false-positive and false-negative results. Moreover, the rapid evolution of viruses, strain-specific host interactions, and the enormous combinatorial search space make exhaustive experimental characterization impractical [18]. Consequently, computational methods have become indispensable for prioritizing high-confidence candidate interactions for subsequent experimental validation and large-scale host–virus interaction analysis [13].

Recent advances in artificial intelligence (AI) and machine learning (ML) have transformed biological interaction prediction by enabling data-driven learning from rapidly expanding biomedical datasets [14]. Among these approaches, graph representation learning and Graph Neural Networks (GNNs) have emerged as powerful computational frameworks for modeling complex biological systems because they explicitly exploit graph topology, relational dependencies, and node features [15]. GNNs have achieved remarkable success across diverse bioinformatics applications, including PPIs prediction, drug–target interactions (DTIs) prediction, disease gene prioritization, biological knowledge graph completion, and systems biology [16]. These advances have established graph learning as a promising paradigm for studying HV–PPIs networks, where the connectivity patterns among host and viral proteins provide rich topological information that can facilitate the discovery of previously uncharacterized HV–PPIs.

Graph representation learning has emerged as an effective paradigm for modeling biological interaction networks by representing proteins as nodes and their interactions as edges within a graph. In this framework, link prediction aims to infer previously unknown interactions by exploiting the underlying network topology and observed interaction patterns rather than relying solely on sequence or structural similarity [17]. Unlike conventional pairwise prediction methods that treat proteins as independent entities, graph-based approaches preserve the interconnected organization of biological systems and enable information propagation across neighboring proteins and functional modules, thereby facilitating the discovery of latent functional relationships. This capability is particularly advantageous for heterogeneous HV–PPIs, where host proteins form dense interaction networks while viral proteins selectively target specific host proteins to manipulate cellular pathways.

The effectiveness of graph-based learning is closely related to the principle of network proximity, which states that proteins involved in similar biological processes or disease mechanisms tend to occupy nearby regions of the interaction network [18]. In host– virus systems, viruses typically perturb interconnected host pathways rather than isolated proteins, implying that biologically meaningful interaction signals may arise from both immediate molecular neighborhoods and distant topological regions of the host interactome [19]. GNNs naturally address this problem through iterative message passing, enabling node representations to incorporate information from neighboring nodes while preserving graph structure [20].

Additionally, most existing computational approaches for HV–PPIs prediction rely on protein sequence information, structural representations, or multimodal feature integration, whereas topology-driven graph learning remains comparatively underexplored. In addition, current GNN-based methods predominantly employ a single graph representation strategy and rarely perform a systematic comparison of local, global, and integrated graph representations within a unified framework. As a result, the individual and complementary contributions of different propagation scales to HV–PPIs prediction remain largely unknown, motivating the development of a dedicated multi-scale graph learning framework. Our previous work, OMNI (Optimized Multiview Network Integration), combined relation-aware graph attention with global propagation to improve representation learning in heterogeneous biomedical networks [21]. Although OMNI demonstrated the effectiveness of integrating multiple graph representations for general biomedical link prediction, it was not specifically designed to investigate how different propagation scales influence HV–PPIs prediction.

Motivated by these challenges, we propose HV–MultiScaleGNN, a multi-scale GNN framework for HV–PPIs prediction. Unlike existing methods that predominantly employ a single graph representation strategy or integrate sequence and structural information, HV– MultiScaleGNN formulates HV–PPIs prediction as a topology-driven heterogeneous link prediction problem and systematically investigates three complementary graph representation strategies: Local view, Global view, and Local+Global view. The Local view employs relation-aware graph attention to capture fine-grained neighborhood interactions, whereas the Global view combines Random Walk with Restart (RWR) and hierarchical attention to model higher-order structural dependencies within the heterogeneous host–virus interaction network. The Local+Global view integrates these complementary representations into a unified embedding space, enabling simultaneous learning from immediate molecular interactions and broader network topology. This unified framework provides a systematic assessment of how different propagation scales influence prediction performance, probability calibration, and ranking capability while offering new insights into the complementary roles of local and global graph representations in HV–PPIs prediction.

The major contributions of this work are summarized as follows:

1. We propose HV–MultiScaleGNN, a topology-driven multi-scale GNN framework that formulates HV–PPIs prediction as a heterogeneous link prediction problem using only heterogeneous network topology.
2. We systematically compare Local, Global, and Local+Global graph representation strategies within a unified computational framework, providing a comprehensive analysis of their individual and complementary contributions to HV–PPIs prediction.
3. We introduce a propagation-guided representation learning framework that integrates relation-aware graph attention with higher-order graph propagation to capture both local neighborhood interactions and global structural dependencies in heterogeneous biological networks.
4. We optimize the general OMNI framework tailored to HV–PPIs.
5. We perform extensive experimental evaluation using discrimination, calibration, ranking, external validation, biological interpretation analyses, structural assessment, GO and KEGG enrichment analysis, demonstrating the effectiveness and interpretability of the proposed framework for topology-driven HV–PPI prediction.

The overall architecture of the proposed HV–MultiScaleGNN framework is illustrated in **Fig. 1**. The framework consists of four major stages: (i) construction of a heterogeneous host–virus interaction network by integrating experimentally validated HV–PPIs with the human PPIs network, (ii) multi-scale representation learning through complementary Local and Global graph representation strategies, (iii) feature fusion to generate unified protein embeddings, and (iv) relation-specific heterogeneous link prediction for identifying previously unknown HV–PPIs.

**Fig. 1.**
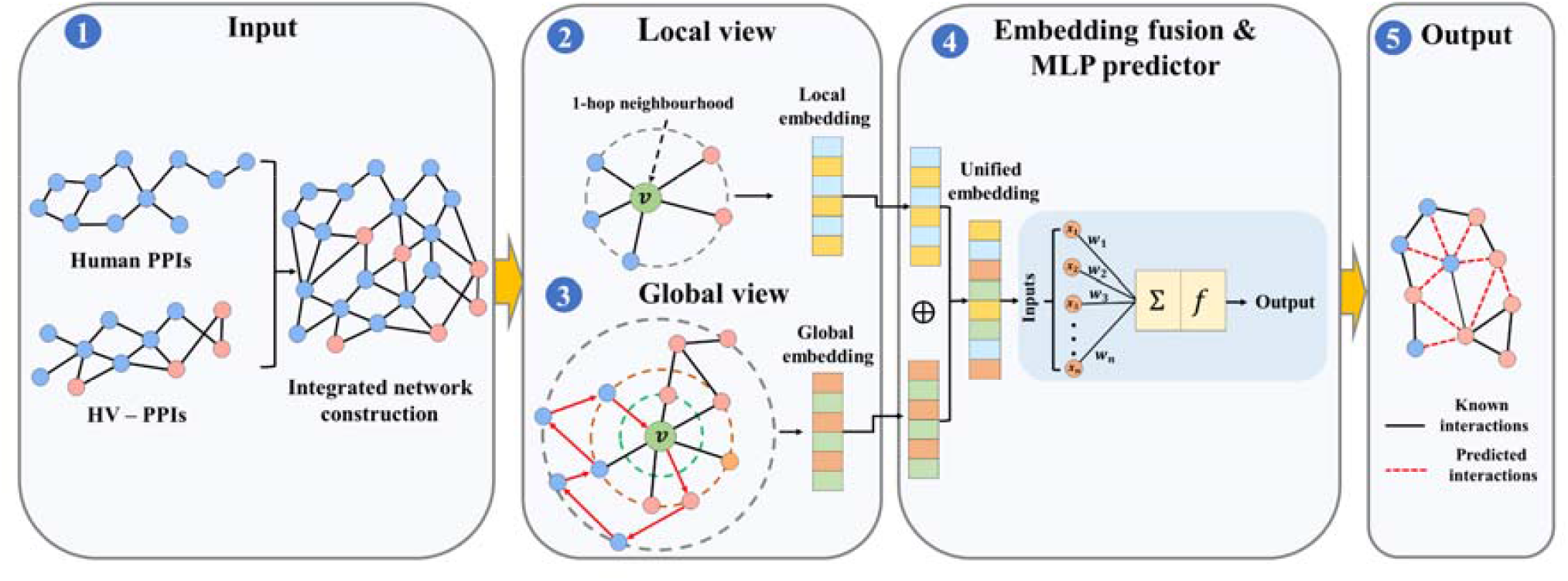
The panorama of the proposed HV-MultiScaleGNN framework for HV–PPIs prediction. The framework constructs a heterogeneous interaction graph by integrating the experimentally validated human PPIs network and HV–PPIs. HV–MultiScaleGNN systematically evaluates three graph representation learning strategies: (i) Local view, (ii) Global view, and (iii) Local+Global view. The Local+Global view integrates the complementary representations learned from the Local and Global branches through a multi-scale feature fusion module to generate unified protein embeddings, which are subsequently processed by a relation-specific MLP decoder for HV–PPIs prediction.

## 2. Related work

The available computational approaches for HV–PPIs prediction have evolved substantially over the past decade. Existing methods can be broadly categorized into four groups: (i) conventional computational and ML approaches, (ii) deep learning (DL) approaches, (iii) GNN-based approaches, and (iv) multi-scale and multi-view graph representation learning approaches.

### 2.1. Conventional computational and ML approaches for HV–PPIs prediction

Computational approaches primarily relied on biological prior knowledge derived from protein sequence similarity, domain–domain interactions, structural information, evolutionary conservation, physicochemical properties, and functional annotations. These handcrafted biological features were combined with conventional ML algorithms, including Support Vector Machines (SVMs), Random Forests (RFs), Naïve Bayes (NB), and ensemble learning methods, to predict potential HV–PPIs [22, 23]. Although these approaches achieved encouraging performance, their dependence on manually engineered features limited their ability to capture the complex nonlinear relationships underlying host–virus interactions and restricted their generalizability across different viruses.

### 2.2. Deep learning for biological interaction prediction

DL has transformed biological interaction prediction by enabling automatic feature learning directly from high-dimensional biological data. Convolutional Neural Networks (CNNs), Recurrent Neural Networks (RNNs), attention mechanisms, transformer architectures, have been successfully applied to protein function prediction, PPIs prediction, DTIs prediction, disease gene prioritization, and related biological applications [24–26]. DeepViral integrated protein sequence representations with host phenotype and functional annotations to improve prediction accuracy and biological interpretability [27]. DeepVHPPI introduced a novel DL framework combining a self-attention-based transformer architecture and a transfer learning training strategy to predict interactions between human proteins and virus proteins that have novel sequence patterns [28]. More recently, DeepHVI, HBFormer, and ECA-PHV combined protein language models with sequence, structural, physicochemical, evolutionary, and functional information to further improve prediction robustness and generalization [29–31]. Despite these advances, most DL-based approaches primarily rely on protein-derived information while making limited use of the topological organization of biological interaction networks.

### 2.3. Graph Neural Networks in biological networks

GNNs have emerged as powerful frameworks for modeling biological systems because proteins, genes, drugs, diseases, and other biomolecules naturally form complex interaction networks. Representative architectures, including Graph Convolutional Networks (GCNs), GraphSAGE, Graph Attention Networks (GATs), Relational Graph Convolutional Networks (R-GCNs), and Heterogeneous Graph Transformers (HGTs), have demonstrated strong performance in PPIs prediction, DTIs prediction, disease gene prioritization, biological knowledge graph completion, and drug repurposing [32–38].

Recent GNN-based studies have further incorporated graph attention mechanisms, heterogeneous graph modeling, and topology-aware message passing to improve biological interaction prediction. DSSGNN-PPI integrates protein structural graphs with sequence information through graph attention [39]. MGPPI combines multiscale graph convolution with an interpretable Gradient Weighted interaction Activation Mapping (Grad-WAM) mechanism to capture complementary structural information [40]. GNNGL-PPI jointly learns global graph representations and local subgraph information for multicategory PPIs prediction [41], whereas CM-PHI integrates heterogeneous microbial interaction networks with multi-hop graph attention for phage–host interaction prediction [42]. Collectively, these studies demonstrate that exploiting graph topology substantially improves biological interaction prediction beyond sequence-only learning.

### 2.4. Multi-scale and multi-view graph representation learning

Multi-scale and multi-view graph representation learning has recently attracted considerable attention because biological networks contain meaningful information at multiple topological scales. Representative techniques include random walk-based propagation, graph diffusion, hierarchical attention, motif-aware aggregation, meta-path reasoning, and multiscale graph convolution. Rather than relying on a single neighborhood aggregation strategy, these methods integrate complementary structural information captured from different propagation depths and graph perspectives to learn more informative node representations. Such approaches have consistently improved representation quality and predictive performance across diverse graph learning applications by jointly exploiting local neighborhood structure and higher-order network topology.

### 2.5 Research gap

Despite significant progress in computational HV–PPIs prediction, several key challenges still persist. First, most sequence-based DL models primarily rely on protein sequence information and often overlook the complex topological organization of biological interaction networks. Second, existing GNN approaches typically focus on either local neighborhood aggregation or higher-order graph propagation without systematically investigating their complementary contributions. Third, although multi-scale and multi-view graph learning approaches have shown promising results, a comprehensive evaluation of Local view, Global view, and integrated Local+Global view representations within a unified heterogeneous host–virus interaction framework remains largely unexplored. Moreover only a few studies have investigated how different propagation scales influence not only predictive discrimination but also probability calibration and ranking capability, both of which are essential for reliable biological link prediction. These limitations motivate the development of HV–MultiScaleGNN, which systematically investigates Local, Global, and Local+Global graph representation strategies within a unified topology-driven graph learning framework based on integrating relation-aware graph attention with global propagation in heterogeneous biomedical networks [21].

## 3. Preliminary

In this section, we formally define the HV–PPIs graph and formulate the HV–PPIs prediction task. We further introduce the multi-scale graph learning concepts that form the foundation of the proposed HV–MultiScaleGNN framework.

### 3.1 Definition 1: Host–Virus interaction graph

Given a heterogeneous graph *G= (V, E)*, the node set *V= V*_*H*_ ∪ *V* _*V*_ consists of host proteins *V*_*H*_ and viral proteins *V*_*V*_ . The edge set *E = E*_*HH*_ ∪ *E*_*HV*_ contains two types of interactions: human PPIs *E*_*HH*_ derived from the human interactome and experimentally validated HV–PPIs *E*_*HV*_. Each node *v* ∈ *V* is associated with a feature vector describing its biological characteristics, while each edge *e* ∈ *E* represents a biological interaction. The objective of HV–PPIs prediction is to identify previously unknown host–virus interactions 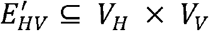.

### 3.2 Definition 2: Local neighborhood graph

For a target node *v*, the local neighborhood graph *N*_*local*_(*v*) consists of its directly connected one-hop neighbors. The local graph captures immediate interaction patterns and preserves fine-grained biological relationships among host and viral proteins.

### 3.3 Definition 3: Global neighborhood graph

For a target node *v*, the global neighborhood graph *N*_*global*_*(v) c*ontains high-order neighbors obtained through RWR sampling on the heterogeneous interaction network. This graph captures long-range dependencies, indirect functional associations, and global topological characteristics of the host–virus interactome.

### 3.4 Definition 4: Multi-Scale graph attention network

The proposed HV–MultiScaleGNN framework systematically investigates three graph representation strategies: Local view, Global view, and Local+Global view. The Local view learns node representations from one-hop neighborhoods using relation-aware graph attention, whereas the Global view captures high-order structural dependencies through RWR and hierarchical attention. The Local+Global view integrates the complementary local and global representations through multi-scale feature fusion.

### 3.5 Definition 5: Host–Virus interaction prediction

Given a host protein *h* ∈ *V*_*H*_ and a viral protein *v* ∈ *V*_*V*_, the prediction task is to estimate the probability of interaction between the protein pair:

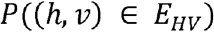

using the protein representations generated by the selected graph representation strategy (Local view, Global view, or Local+Global view).

## 4. Methodology

### 4.1. HV–MultiScaleGNN model architecture

Let *G= (V, E)* denote the combined homogenous human PPIs graph and heterogeneous HV–PPIs graph, where *V* = {*v*_*i*_} represents the set of host and viral proteins, and *E* = {(*v*_*i*_, *r, v*_*j*_ )} denotes the set of hosts–virus interactions. Here, *r* corresponds to the interaction type, including human PPIs and HV–PPIs. The objective of HV–MultiScaleGNN is to estimate the probability of a candidate host–virus interaction *e*_*ij*_ = (*v*_*i*_, *r, v*_*j*_). We propose HV–MultiScaleGNN, a multi-scale GNN framework that formulates HV–PPIs prediction as a topology-driven heterogeneous link prediction problem using an encoder– decoder architecture. The encoder systematically investigates three complementary graph representation strategies, namely Local view, Global view, and Local+Global view. In the Local view, relation-aware graph attention aggregates information from one-hop neighbors to learn fine-grained local protein representations. In the Global view, RWR-based neighborhood sampling together with hierarchical attention captures high-order topological dependencies and long-range structural information within the heterogeneous interaction network. The Local+Global view integrates the complementary local and global representations through a multi-scale feature fusion module with type-specific projection to generate unified protein embeddings. The protein representations generated by the selected representation strategy are subsequently provided to a relation-specific interaction prediction module, which estimates the likelihood of interaction between candidate host and viral protein pairs. Consequently, HV–MultiScaleGNN enables a systematic comparison of local, global, and integrated multi-scale representations within a unified learning framework while sharing the same heterogeneous graph construction, decoder architecture, optimization strategy, and evaluation protocol. The overall computational workflow of HV– MultiScaleGNN is summarized in **Algorithm 1**.

#### Algorithm 1

HV-MultiScaleGNN framework for HV–PPIs Predictions

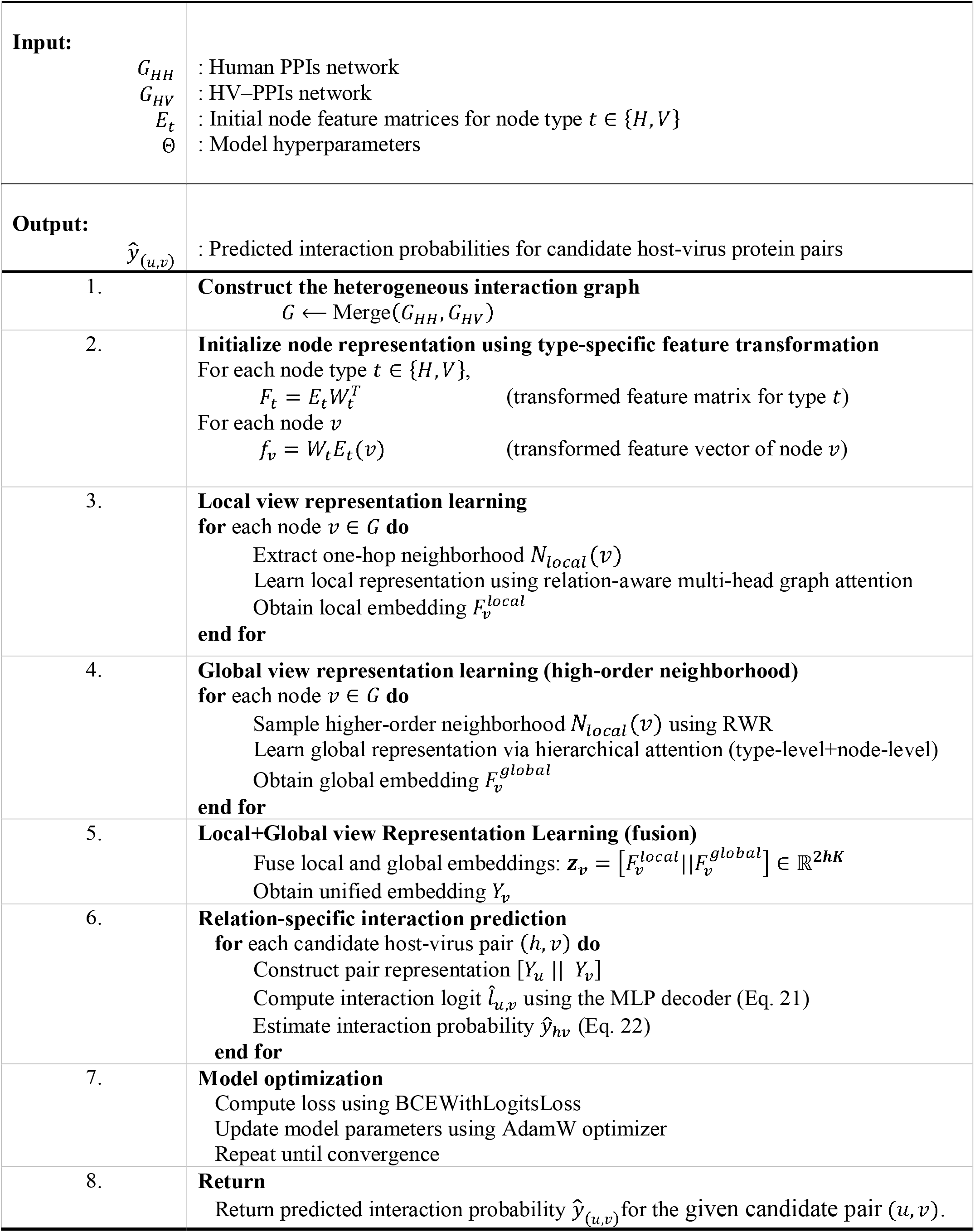

### 4.2. Data collections and pre-processing

Experimentally validated human PPIs and HV–PPIs were collected from publicly available biological databases to construct the heterogeneous HV–PPIs network. Human PPIs were integrated from multiple curated resources, including BioGRID, the human PPIs compiled by Gysi et al., the SNAP human PPIs dataset, and the human PPIs reported by Menche et al., to obtain a comprehensive representation of the human interactome [43–45, 18]. The compiled human interactome consists of 1,048,577 PPIs (edges) and 24,482 unique human proteins, providing a comprehensive network for modeling host cellular interactions. Experimentally validated HV–PPIs were obtained from the Human Virus Interaction Database (HVIDB), a comprehensive repository of experimentally validated human–virus interactions [46]. HVIDB contained 48,643 experimentally verified HV–PPIs spanning 35 virus families, which served as the initial host–virus interaction dataset for this study as on 08^th^ April 2026. The summary statistics of the human PPIs datasets and the host–virus interaction dataset both before and after the preprocessing procedure are presented in **Table 1**.

**Table 1.** Summary statistics of the human PPIs and the HV-PPIs dataset used for constructing the heterogeneous interaction network.

| **Dataset** | **Original Interactions** | **Duplicate Interactions Removed** | **Unique Interactions** | **Unique Proteins (Nodes)** |
|---|---|---|---|---|
| Human PPIs (BioGRID) [61] | 1,220,590 | – | 1,220,590 | 20,262 |
| Human PPIs [62] | 327,924 | – | 327,924 | 18,505 |
| Human PPIs (SNAP) [63] | 342,353 | – | 342,353 | 21,557 |
| Human PPIs [28] | 141,296 | – | 141,296 | 13,460 |
| Compiled Human Interactome (after merging all human PPIs datasets and removing duplicate interactions) | 2,032,163 | 983,586 | 1,048,577 | 24,482 |
| HV-PPIs (HVIDB Original) | 48,643 | – | 48,643 | 1,165 viral proteins |
| HV-PPIs (Expanded by UniProt Mapping) | 49,440 | 18,316 | 31,124 | 1,165 viral proteins |

Prior to network construction, the collected datasets underwent several preprocessing steps to improve data consistency and remove redundancy. In the HVIDB dataset, certain interaction records included multiple human Gene IDs mapped to a single UniProt accession. These entries, which were represented as semicolon-separated Gene IDs (e.g., 121504; 554313; 8294; 8359; etc.) were expanded into individual interaction records to create a unique mapping between individual human genes and their corresponding viral protein. As a result of this preprocessing procedure, the total number of interaction records rose from 48,643 to 49,440. The redundant database annotations were subsequently removed. After removing the duplicate records, the dataset contained 31,124 unique experimentally validated HV–PPIs. Furthermore, viral proteins sharing identical protein names but originating from different viral strains, isolates, geographical variants, or sampling locations were consolidated into a single viral protein entity, resulting in 1,165 unique viral proteins used for heterogeneous network construction. This normalization procedure reduced annotation redundancy while preserving biologically equivalent protein functions across different viral variants. The resulting processed datasets provide a high-quality and non-redundant collection of human PPIs and HV–PPIs, which were subsequently used to construct the heterogeneous interaction network. The processed HV–PPIs dataset and the compiled human interactome dataset used in this study are provided in the **Supplementary file S1 and S2**.

### 4.3. Network construction

The preprocessed human PPIs and HV–PPIs dataset were integrated to construct a heterogeneous host–virus interaction graph for representation learning and interaction prediction. In the proposed graph, all human and viral proteins were represented as graph nodes, whereas experimentally validated PPIs were represented as edges.

The heterogeneous graph consists of two biologically distinct interaction types. The first comprises human PPIs, which describe molecular interactions within the human interactome. The second comprises HV–PPIs, representing experimentally validated interactions between human and viral proteins. Integrating these complementary interaction types enables the proposed framework to simultaneously capture intra-species functional associations and inter-species infection-related interactions within a unified heterogeneous graph.

Following integration, the whole network (heterogeneous) consisted of 1,079,699 human PPIs (edges) comprising of 24,499 unique human proteins and 1,165 unique viral proteins. The combined heterogeneous graph therefore contains two node types (human proteins and viral proteins) and providing a biologically comprehensive representation of the host–virus interaction landscape. The constructed heterogeneous graph serves as the input for the new HV–PPIs prediction.

### 4.4. Feature transformation

Direct aggregation of heterogeneous node features without prior alignment may introduce representation bias and hinder effective information propagation. Therefore, a type-specific feature transformation layer is employed to project all node representations into a common latent space before neighborhood aggregation [47, 36]. Let *T* = {*H,V*} denote the set of node types, where *H* and *V* represent host proteins and viral proteins, respectively. For each node type *t* ∈ *T*, let *V*_*t*_ = {*v* | *v* belongs to node type *t*} denote the corresponding set of nodes. A learnable embedding matrix is assigned to each node type: 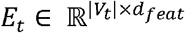, where *d*_*feat*_ denotes the initial feature dimension and |*V*_*t*_| represents the number of nodes belonging to type *t*. To map heterogeneous node representations into a shared hidden space, a type-specific projection matrix 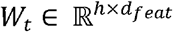 is learned for each node type, where *h* denotes the hidden embedding dimension. For a node *v* ∈ *V*_*t*_, the transformed feature representation is computed as

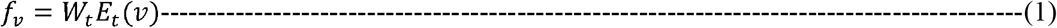

where *E*_*t*_(*v*) denotes the embedding vector of node *v*, and *f*_*v*_ ∈ ℝ^*h*^ represents the projected feature in the common latent space. The transformed representations enable host and viral proteins to be embedded in a unified feature space while preserving type-specific biological characteristics. This alignment facilitates information exchange across heterogeneous interactions during subsequent graph attention operations. For computational efficiency, the transformed feature matrix of each node type is precomputed as

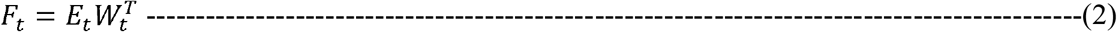

where 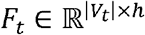 denotes the transformed feature matrix for all nodes belonging to node type *t*. The resulting feature matrices are cached and reused during both local and global neighborhood aggregation. This strategy avoids repeated projection operations over the entire graph and significantly reduces computational overhead during training. The transformed node representations obtained from Eq. (1) and (2) serve as the input features for the subsequent Local view and Global view representation learning strategies.

### 4.5. Local neighborhood representation learning

The local view representation learning strategy is designed to capture fine-grained interaction patterns from the immediate one-hop neighborhood of each protein in the heterogeneous HV–PPIs network. To effectively model heterogeneous host–virus interactions, Local view employs a relation-aware multi-head graph attention mechanism that sequentially performs relation-specific feature projection, relation-aware attention computation, attention normalization, weighted neighborhood aggregation, multi-head and relation-level feature integration, and residual feature refinement [34, 47]. The detailed architecture of the Local view representation learning strategy within the proposed HV– MultiScaleGNN framework is illustrated in **Fig. 2**. Collectively, these computational operations enable the Local view encoder to learn discriminative protein representations by capturing fine-grained biological interaction patterns from the immediate one-hop neighborhood of each protein.

**Fig. 2.**
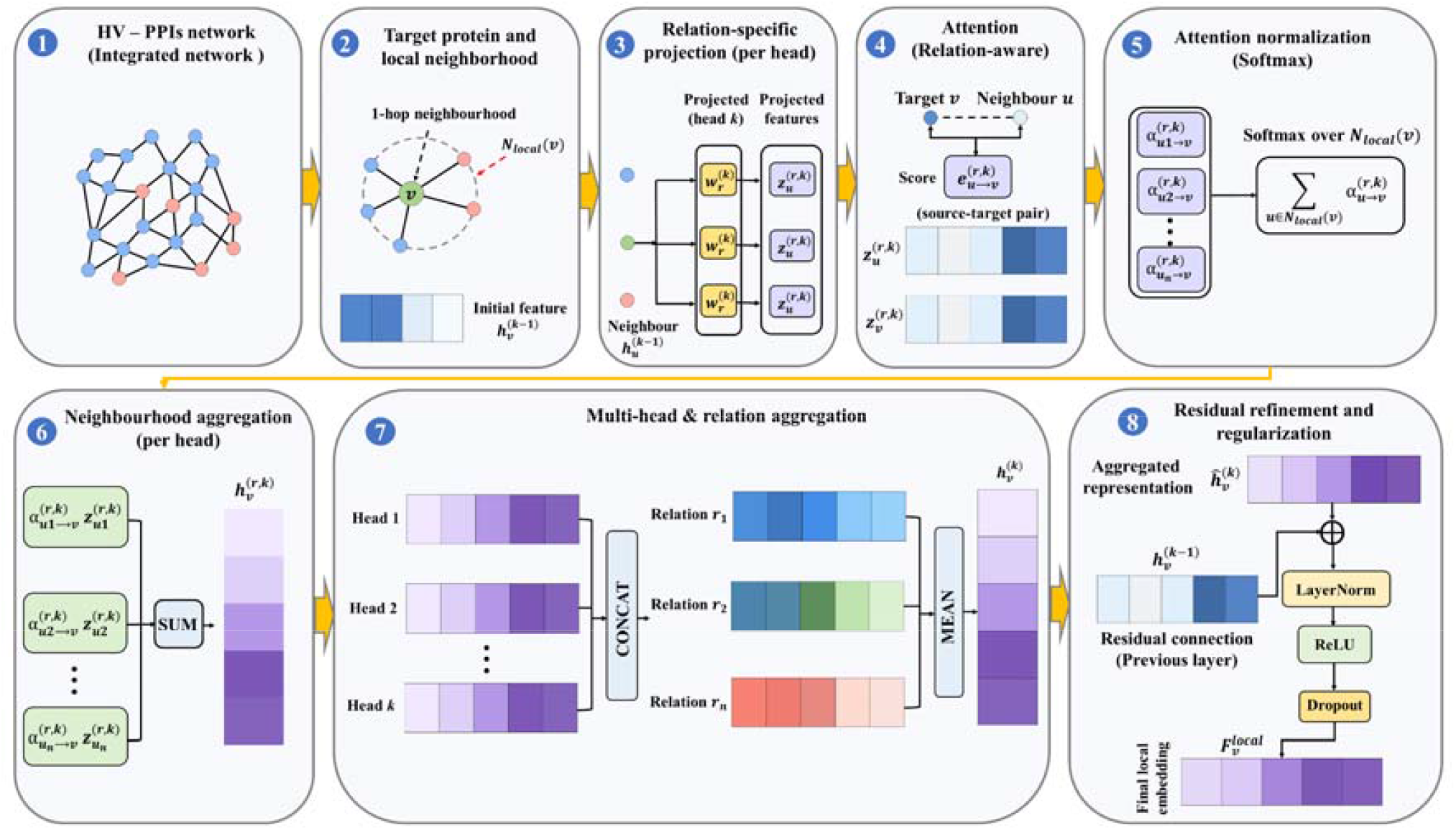
Detailed architecture of the Local view representation learning strategy in HV– MultiScaleGNN. Starting from the heterogeneous host–virus interaction network, the Local view learns fine-grained protein representations through a relation-aware multi-head graph attention mechanism. The computation sequentially performs (1) input heterogeneous HV– PPIs network, (2) one-hop neighborhood extraction around the target protein, (3) relation-specific feature projection for each attention head, (4) relation-aware attention score computation, (5) attention normalization using softmax, (6) weighted neighborhood aggregation for each attention head, (7) multi-head and relation-level feature integration, and (8) residual feature refinement with LayerNorm, ReLU, and dropout to generate the final local node embedding.

#### 4.5.1. Relation-specific projection

For a target protein *v*, let 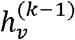 represent its feature embedding at layer *k −* 1. For each neighbor *u* connected by a specific relation *r*, we first project the node features into a relation-specific space:

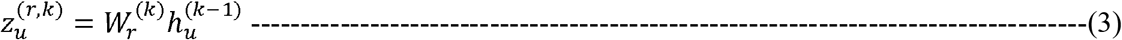

where 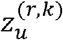 denotes the relation-specific projected representation of neighboring protein *u*, 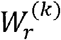 is the learnable projection matrix associated with relation *r*, and 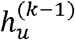 represents the feature embedding of neighboring protein *u* from the previous graph layer. By learning relation-specific projections, the proposed encoder maps neighboring proteins into distinct latent subspaces according to their interaction types, enabling subsequent attention operations to capture heterogeneous biological relationships more effectively.

#### 4.5.2. Relation-aware attention

We calculate the importance of a neighbor *u* to the target protein *v* using a shared attention mechanism. The unnormalized attention coefficient 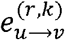 is computed as:

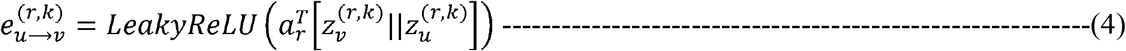

where 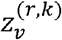 and 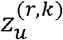 denote the relation-specific representations of the target and neighboring proteins, respectively; *a*_*r*_ is a learnable attention vector associated with relation *r*; and || denotes vector concatenation. The computed attention score quantifies the relative importance of neighboring protein *u* in updating the representation of the target protein *v*. The attention coefficients are subsequently normalized using the softmax function.

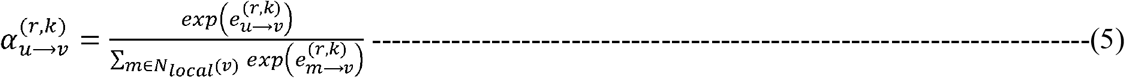

where *N*_*locaol*_(*v*)denotes the set of neighboring proteins connected to target protein *v* through relation *r*, 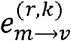 represents the unnormalized attention score between neighboring protein *m* and target protein *v* computing using Eq. (4), and 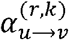 represents the normalized attention coefficient assigned to neighboring protein *u*. The softmax normalization ensures that the attention coefficients are comparable across all neighboring proteins and sum to one, allowing biologically informative interactions to receive larger weights during neighborhood aggregation.

#### 4.5.3. Aggregation and Multi-head integration

The neighborhood information associated with relation *r* is aggregated using the normalized attention coefficients to obtain a relation-specific protein representation.

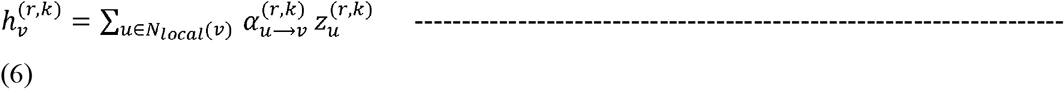

where 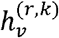 denotes the aggregated relation-specific representation of the target protein *v*, 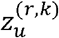 represents the projected feature vector of neighboring protein *u*, and 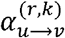 is the normalized attention coefficient. The weighted aggregation enables the Local view encoder to integrate information from neighboring proteins while assigning greater importance to biologically informative interactions according to their learned attention weights.

The Local view representation strategy employs multi-head attention to improve representation capacity and stabilize model training. The relation-specific representations generated by different attention heads are concatenated and projected into a unified embedding space according to

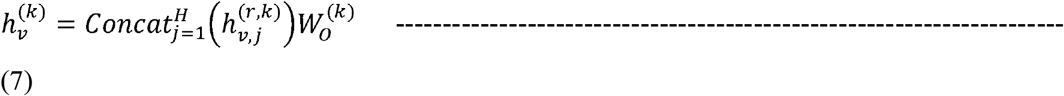

where *H* denotes the number of attention heads, Concat (·) represents feature concatenation across all attention heads, and 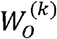 is the learnable output projection matrix. The resulting multi-head representation is subsequently integrated across relations to obtain the final local protein representation. Multi-head attention enables the encoder to capture complementary biological interaction patterns from multiple representation subspaces, thereby improving the robustness and expressiveness of Local view representation learning.

#### 4.5.4. Residual feature refinement

A residual connection followed by layer normalization, ReLU activation, and dropout regularization is employed to preserve the original node information and facilitate stable gradient propagation during training,

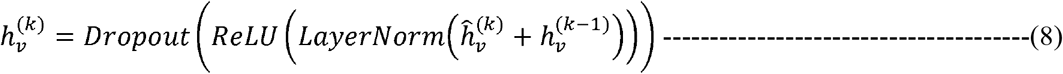

where 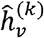 denotes the aggregated neighborhood representation prior to residual refinement. LayerNorm, ReLU, and Dropout denote layer normalization, rectified linear unit activation, and dropout regularization, respectively. The residual pathway preserves the intrinsic characteristics of the target protein while facilitating stable gradient propagation and reducing information loss during iterative message passing. The refined embedding obtained after residual refinement constitutes the final local representation of node *v*, denoted as

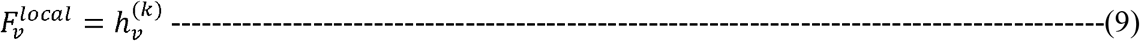

### 4.6. Global neighborhood representation learning

The global view representation learning strategy is designed to capture high-order structural dependencies that extend beyond the immediate neighbourhood of each protein in the heterogeneous host–virus interaction network. Within HV–MultiScaleGNN, the Global view combines RWR-based neighborhood sampling with hierarchical attention to learn complementary global protein representations. The Global view first identifies informative high-order neighbors through RWR sampling and subsequently employs type-level and node-level attention mechanisms to selectively aggregate biologically relevant information. Finally, the aggregated global representation is projected into the latent embedding space, producing a complementary global protein representation that captures long-range topological information. The detailed architecture of the Global view representation learning strategy within the proposed HV–MultiScaleGNN framework is illustrated in **Fig. 3**. The computation sequentially performs RWR-based neighborhood sampling, hierarchical attention (type-level and node-level), attention-guided global neighborhood aggregation, and residual feature projection to generate the final global protein representation.

**Fig. 3.**
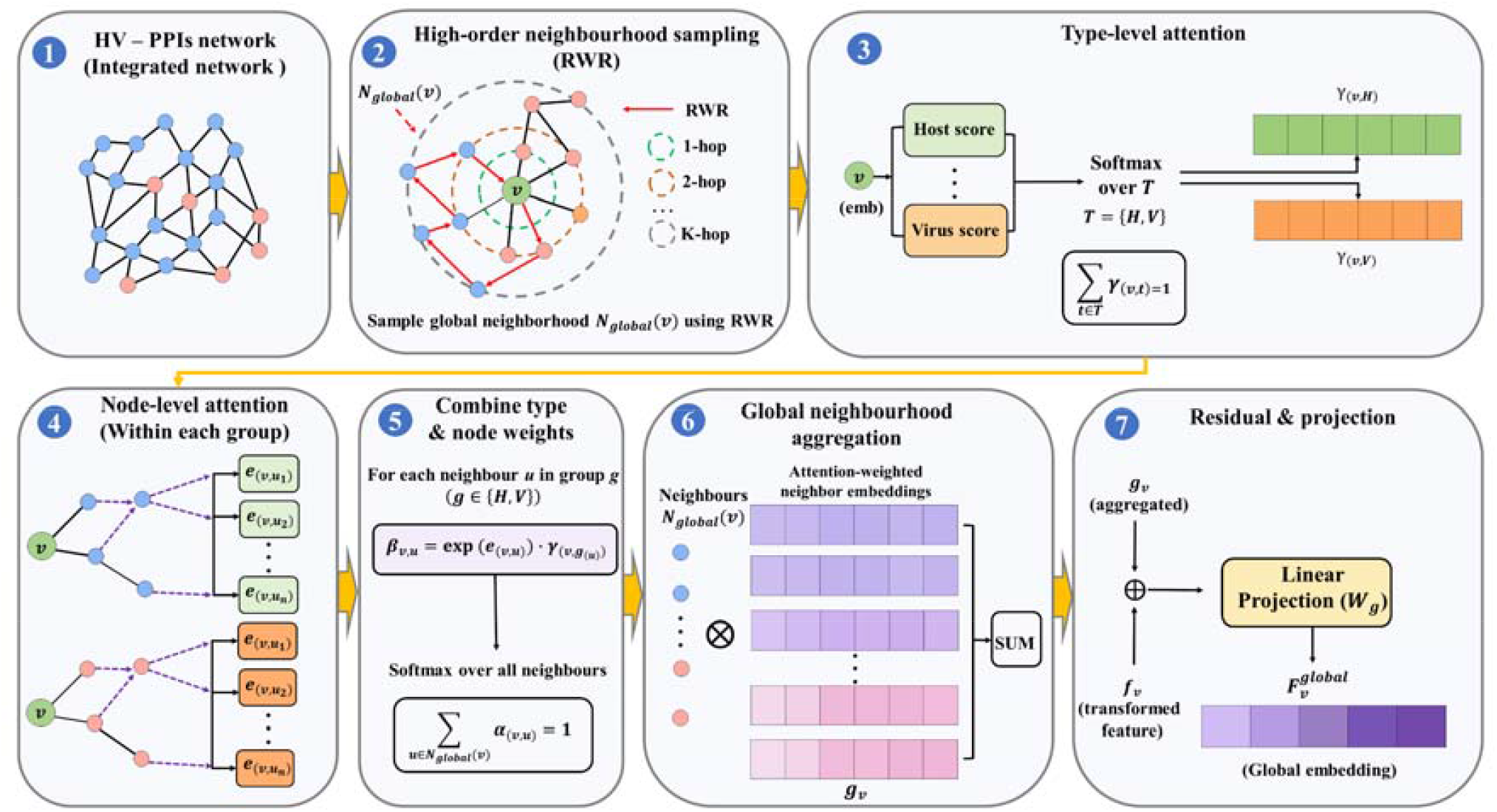
Detailed architecture of the Global view representation learning strategy in HV– MultiScaleGNN. Starting from the constructed heterogeneous host–virus interaction network, the Global view learns high-order protein representations through propagation-guided neighborhood exploration and hierarchical attention. The computation sequentially performs (1) constructed heterogeneous HV–PPIs network as input, (2) Random Walk with Restart (RWR)-based high-order neighborhood sampling, (3) type-level attention to estimate the relative importance of different node types, (4) node-level attention score computation within each node type, (5) combination of type-level and node-level attention scores followed by softmax normalization to obtain the final attention coefficients, (6) attention-weighted global neighborhood aggregation, and (7) residual feature integration with linear projection to generate the final global node embedding.

#### 4.6.1. Random Walk with Restart (RWR)

Unlike local neighborhood aggregation, which focuses exclusively on directly connected proteins, the global branch explores higher-order network topology through the RWR algorithm [48]. RWR iteratively alternates between traversing adjacent nodes and returning to the initial (source) protein, allowing the model to identify distant proteins that are functionally associated even in the absence of direct interactions. For a target protein *v*, the global neighborhood is defined as

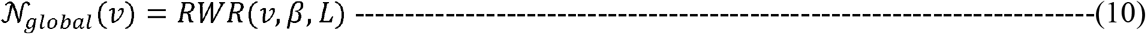

where *β* denotes the restart probability, which controls the trade-off between local exploration and global network traversal, and *L* represents the maximum length of the random-walk. The resulting neighborhood includes higher-order host and viral proteins, thereby providing complementary structural information that extends beyond immediate biological interactions.

#### 4.6.2 Type-level attention mechanism

The sampled global neighborhood consists of proteins belonging to different biological node types, each contributing differently to the representation of the target protein. To account for this heterogeneity, the Global view representation strategy in HV– MultiScaleGNN employs a type-level attention mechanism that automatically learns the relative importance of host and viral proteins during global representation learning.

Let *T* = {*H,V*} denote the set of node types, where *H* and *V* represent host and viral proteins, respectively. For a target protein *v*, let *m*_*v,t*_ denote the aggregated representation of neighboring proteins belonging to node type *t*. The importance score of each node type is computed as

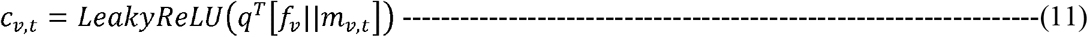

where *f*_*v*_ denotes the transformed feature representation of the target protein, *q i*s a learnable attention vector, and || represents vector concatenation.

The type-level attention scores are normalized using the softmax function

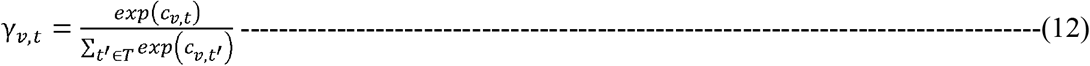

where *γ*_*v,t*_ denotes the normalized attention weight assigned to node type *t*. The resulting attention coefficients enable the Global view encoder to emphasize biologically informative node types while suppressing less relevant node types during global neighborhood aggregation.

#### 4.6.3 Node-level attention score estimation

Following type-level attention, the importance of each sampled high-order neighboring protein is estimated through a node-level attention mechanism. Rather than assigning equal importance to all sampled neighbors, the Global view representation strategy learns adaptive attention weights according to the feature similarity between the target protein and each neighboring protein. For a target protein *v* and one of its sampled neighbors *u*, the unnormalized attention score is computed as

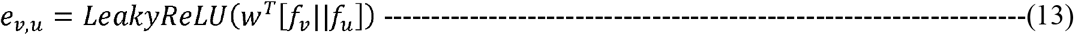

where *f*_*v*_ and *f*_*u*_ denote the feature representations of the target and neighboring proteins, respectively, *w* is a trainable attention vector, and [·||·] denotes vector concatenation. The LeakyReLU activation introduces nonlinearity into the attention computation.

Since the Global view representation strategy employs a hierarchical attention mechanism, the node-level attention score is further combined with the corresponding type-level attention coefficient. The combined attention coefficient is computed as

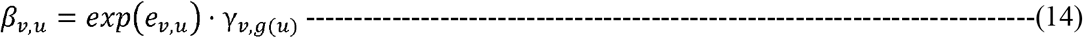

where *γ*_*v,g(u)*_ represents type-level attention weight assigned to the node-type group containing neighboring protein *u*, and *g*(*u*) denotes the corresponding node type associated with neighbor *u*.

To obtain normalized attention coefficients, the combined attention scores are normalized across all sampled high-order neighbors using the Softmax function

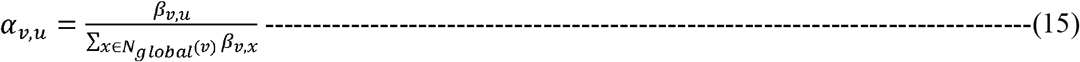

where *N*_*global*_ (*v*)denotes the set of high-order neighbors sampled for target protein *v*. The normalized attention coefficients satisfy

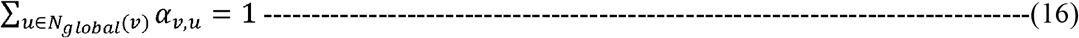

thereby allowing the attention coefficients to be interpreted as probability-like importance weights during global neighborhood aggregation.

By jointly considering node-level feature similarity and type-level importance, the proposed hierarchical attention mechanism enables the Global view representation strategy to effectively distinguish informative high-order biological interactions from less relevant network connections. Consequently, the learned global representations capture long-range structural dependencies while reducing the influence of noisy neighbors, resulting in more discriminative protein embeddings for HV–PPIs prediction.

#### 4.6.4 Global neighborhood aggregation and feature projection

Within the Global view representation strategy, the normalized attention coefficients are used to aggregate information from the sampled high-order neighbors, thereby constructing a complementary global representation for each protein. Unlike the Local view, which captures immediate biological interactions, the Global view aggregates indirect functional associations distributed throughout the heterogeneous interaction network. The aggregated global representation is computed as

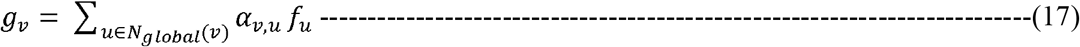

where *g*_*v*_ denotes the aggregated global representation of protein *v*; *α*_*v, u*_ is the normalized attention coefficient; *f*_*u*_ represents the transformed feature vector of neighboring protein *u*; and *N*_*global*_ (*v*)denotes the set of high-order neighbors obtained through RWR sampling.

A residual self-connection is incorporated by combining the aggregated neighborhood representation with the original transformed feature to preserve the intrinsic characteristics of the target protein during message propagation. The combined representation is then projected into the final global embedding space using a learnable linear transformation

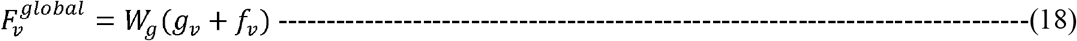

where *W*_*g*_ is a learnable projection matrix that maps the combined global neighborhood representation into the shared latent embedding space, *g*_*v*_ denotes the aggregated global neighborhood representation obtained from Eq. (17), and *f*_*v*_ denotes the transformed node feature vector obtained from the feature transformation layer.

The resulting embedding constitutes the final Global view representation. Within the proposed HV–MultiScaleGNN framework, this representation can be directly utilized for interaction prediction in the Global view configuration or integrated with the Local view representation through the multi-scale feature fusion module to generate unified protein embeddings for the Local+Global view configuration.

### 4.7. Multi-scale feature fusion

The local and global representation learning strategies capture complementary structural information from the heterogeneous host–virus interaction network. The Local+Global view representation strategy integrates local neighborhood information with higher-order topological context to generate biologically informative unified node embeddings. The fusion process consists of three sequential stages: (i) feature concatenation, (ii) type-specific linear projection, and (iii) unified node representation learning.

#### 4.7.1 Feature concatenation

The first stage combines the embeddings obtained from the local and global view representation learning strategy using vector concatenation. This operation retains the complete information learned from both the Local and Global view representation strategies without discarding any structural characteristics. For each protein node *v*, the concatenated multi-scale feature vector is defined as

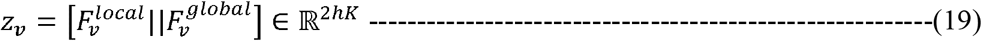

where 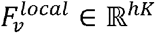 represents the local node representation; 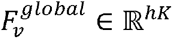 corresponds to the global node representation; and || denotes the vector concatenation operator.

The concatenated representation maintains the complementary information captured from both neighborhood scales.

#### 4.7.2 Type-specific linear projection

Although the concatenated feature vector integrates complementary structural information from both graph views, host proteins and viral proteins exhibit distinct biological characteristics. To effectively integrate and produce combined representations, the Local+Global view representation strategy employs a type-specific linear projection to transform the concatenated representation into a unified latent embedding space while preserving node-type-dependent information. For each protein node *v* belonging to the node type *t* ∈ {*H,V*}, where *H* and *V* denote host proteins and viral proteins, respectively, the projected representation is computed as

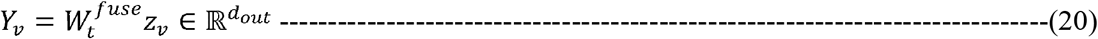

where *z*_*v*_ ∈ ℝ^2*hK*^is the concatenated multi-scale feature vector; 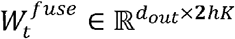 denotes the learnable projection matrix corresponding to node type *t;* and *d*_*out*_ represents the dimensionality of the final embedding space.

The projection layer learns independent transformation parameters for host and viral proteins, enabling the fused Local+Global representations to preserve node-type-specific biological characteristics while projecting both protein types into a common latent embedding space for subsequent interaction prediction. For the Local+Global view representation strategy, *Y*_*v*_ denotes the unified protein embedding obtained through multi-scale feature fusion. During relation-specific interaction prediction, *Y*_*v*_ generally denotes the final protein embedding supplied to the MLP decoder. Accordingly, *Y*_*v*_ corresponds to 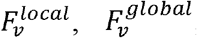, or the fused embedding generated by Eq. (20), depending on whether the Local view, Global view, or Local+Global view representation strategy is employed.

#### 4.7.3 Unified node representation

Following the type-specific linear projection, the resulting embedding *Y*_*v*_ serves as the unified multi-scale protein representation that integrates complementary structural information from the Local view and Global view. By directly projecting the concatenated multi-scale feature vector, the framework enables the neural network to implicitly learn the optimal weighting and integration of local and global signals without imposing artificial scalar constraints.

These final embeddings simultaneously preserve fine-grained, first-order molecular interactions along with higher-order network topology, providing robust and discriminative protein representations for the subsequent relation-specific interaction prediction module. The detailed computational architecture of the Local+Global view representation strategy within the proposed HV–MultiScaleGNN framework is illustrated in **Fig. 4**. Starting from the local and global protein representations learned by the corresponding representation strategies, the Local+Global view performs feature concatenation, type-specific linear projection, and unified protein representation learning before relation-specific interaction prediction.

**Fig. 4.**
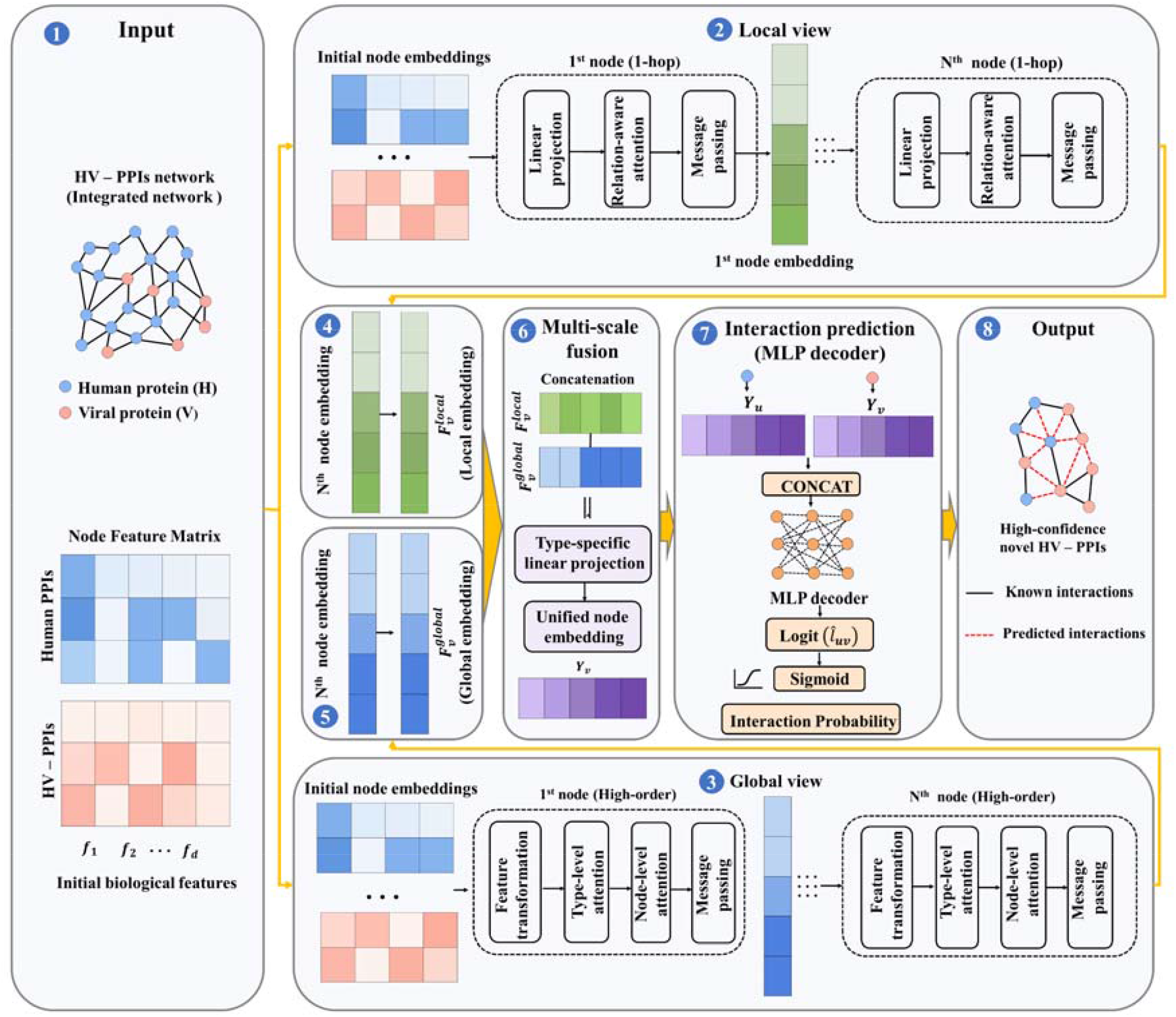
Detailed computational architecture of the Local+Global view representation strategy in HV–MultiScaleGNN. The architecture consists of (1) the input heterogeneous HV–PPIs network and initial node feature representations, (2) the Local view representation learning module, which learns fine-grained one-hop neighborhood representations through relation-aware graph attention, (3) the Global view representation learning module, which captures higher-order topological dependencies through propagation-guided neighborhood exploration and hierarchical attention, (4) extraction of the final local node embeddings, (5) extraction of the final global node embeddings, (6) multi-scale feature fusion with type-specific linear projection to generate unified node embeddings, (7) relation-specific MLP decoder for HV–PPIs prediction, and (8) prediction of high-confidence HV–PPIs.

### 4.9. Model training and interaction prediction

The final protein representations learned by the proposed HV–MultiScaleGNN framework are utilized to predict the likelihood of interactions between host and viral proteins. These protein embeddings are obtained from one of the three graph representation strategies investigated in this study, namely Local view, Global view, or Local+Global view, and subsequently serve as the input to the relation-specific interaction prediction module. Although the ultimate objective of the proposed HV–MultiScaleGNN framework is the identification of previously unknown HV–PPIs, the framework employs a multi-task learning strategy during training. The protein embeddings learned by the selected graph representation strategy are provided as input to the relation-specific MLP-based interaction predictor. For the Local+Global view, these embeddings are obtained through multi-scale feature fusion, whereas for the Local view and Global view, the corresponding branch representations are used directly. These predictors simultaneously estimate the probability of interaction for both candidate HV–PPIs pairs and auxiliary human PPIs.

By jointly optimizing the reconstruction of both interaction types, the model leverages the rich intra-species connectivity of the human interactome to learn biologically more robust and informative protein embeddings. The relation-specific interaction predictors are trained using experimentally validated interactions as positive samples, paired with randomly generated non-interacting protein pairs as negative samples for each respective relation. The model parameters are optimized by minimizing the Binary Cross-Entropy (BCE) loss, enabling the framework to effectively distinguish interacting from non-interacting protein pairs. The selected graph representation strategy and the relation-specific interaction prediction module are jointly optimized in an end-to-end manner, allowing the learned protein representations to be continuously refined according to the interaction prediction objective. By leveraging the protein embeddings learned by the selected graph representation strategy, the proposed interaction prediction module effectively utilizes local interaction patterns, global network topology, or their complementary integration to facilitate accurate identification of previously unknown HV–PPIs.

### 4.10. Relation-specific interaction prediction

The final protein representations generated by the selected graph representation strategy are utilized to estimate the interaction strength between candidate host–virus protein pairs. Given a host protein *u* and a viral protein *v*, HV–MultiScaleGNN predicts the interaction likelihood by concatenating their corresponding protein embeddings and passing the resulting feature vector to MLP. The MLP transforms the combined protein representation into a scalar logit, which reflects the confidence that the corresponding host– virus protein pair forms a biological interaction. The interaction logit is computed as

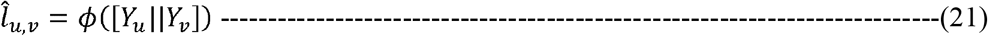

where *Y*_*u*_ and *Y*_*v*_ denote the final protein embeddings generated by the selected graph representation strategy. For the Local+Global view, these correspond to the unified protein embeddings obtained through multi-scale feature fusion, whereas for the Local view and Global view, they correspond to the respective branch-specific embeddings of the host protein and viral protein respectively, || represents the vector concatenation operation, and *ϕ* (·) denotes the two-layer MLP employed as the interaction prediction module.

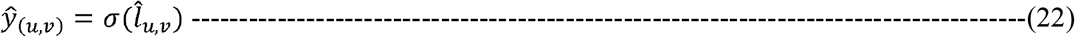

where *σ*(⋅) denotes the sigmoid activation function and *ŷ*_*(u, v)*_ ∈ [0,1] represents the predicted interaction probability for the candidate host–virus protein pair.

The predicted logit summarizes the interaction evidence extracted from both local neighborhood information and global network topology. Higher logit values indicate a greater likelihood that the corresponding host and viral proteins interact. During training, these logits are subsequently transformed into interaction probabilities through the sigmoid activation function and optimized using the BCE objective described in the following section.

### 4.11. Binary Cross-Entropy loss

The interaction prediction module is trained in a supervised manner using a BCE loss with logits to optimize the proposed HV–MultiScaleGNN framework. During training, both experimentally validated HV–PPIs and known human PPIs are treated as positive samples for their respective relation types. Corresponding negative samples are generated by randomly sampling non-interacting protein pairs within the same relation type. This multi-task learning strategy enables the model to simultaneously analyse the human interactome and predict host–virus interactions by averaging the loss across all relation types. The overall training objective is defined as:

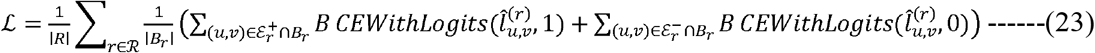

where ℛ denotes the set of all relation types (i.e., human–human and host–virus interactions), |*R*| is the total number of relation types, and | *B*_*r*_ | denotes the mini-batch size for relation *r*. 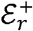 and 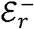 represent the sets of experimentally validated positive interactions and sampled negative interactions under relation *r*, respectively, and 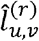 is the predicted interaction logit generated by the relation-specific MLP decoder.

The BCE loss with logits combines the sigmoid activation and cross-entropy objective into a numerically stable optimization function, thereby improving training stability and convergence during deep neural network optimization. During optimization, the model is encouraged to assign higher prediction scores to interacting HV–PPIs pairs while suppressing the scores of non-interacting protein pairs. Consequently, the learned protein embeddings become increasingly discriminative, enabling more accurate identification of previously unknown HV–PPIs.

The encoder and relation-specific interaction prediction module are optimized jointly through end-to-end backpropagation, allowing the multi-scale protein representations to be continuously refined according to the interaction prediction objective. This joint optimization enables HV–MultiScaleGNN to learn biologically meaningful protein representations by exploiting local interaction patterns, global network topology, or their complementary integration, depending on the selected graph representation strategy resulting in robust and biologically meaningful protein representations for HV–PPIs prediction.

### 4.12. Optimization and negative sampling

The proposed HV–MultiScaleGNN framework is trained in an end-to-end manner using the AdamW optimizer, which combines adaptive gradient optimization with decoupled weight decay regularization to improve convergence stability and reduce overfitting during model training [49]. The optimizer iteratively updates all trainable parameters, including the feature transformation layers, attention mechanisms, multi-scale feature fusion module, and relation-aware interaction prediction layer.

Since experimentally verified HV–PPIs constitute only a small fraction of all possible host–virus protein pairs, the interaction prediction problem is formulated as a binary classification task comprising both positive and negative samples. Positive samples represent to experimentally validated HV–PPIs, whereas negative samples are generated by randomly sampling host–virus protein pairs that are not observed in the interaction dataset. This negative sampling strategy allows the model to more effectively distinguish interacting from non-interacting protein pairs while alleviating the severe class imbalance commonly encountered in biological interaction networks [50].

During each training iteration, mini-batches are constructed by combining observed positive interactions with an equal number of sampled negative interactions. The generated negative samples are dynamically refreshed throughout training to expose the model to diverse non-interacting protein pairs and improve its generalization capability. This strategy reduces sampling bias and encourages the framework to learn more discriminative representations of host and viral proteins.

The network parameters are optimized by minimizing the BCE loss described in the previous section. Gradients are computed through backpropagation, and all components of the proposed architecture are jointly optimized in an end-to-end fashion. Overall, the combination of adaptive optimization, dynamic negative sampling, and end-to-end parameter learning enables HV–MultiScaleGNN to effectively learn robust multi-scale protein representations while accurately identifying previously unknown HV–PPIs.

### 4.13. Hyperparameter selection and sensitivity analysis

The predictive performance of GNN is strongly influenced by the selection of hyperparameters, including the embedding dimension, number of attention heads, hidden layer size, learning rate, dropout rate, and weight decay. Rather than relying on manually selected values, HV–MultiScaleGNN employs an automated hyperparameter optimization strategy to identify configurations that maximize predictive performance while maintaining stable model convergence. The hyperparameter search was conducted using the Optuna optimization framework, which efficiently explores the search space through sequential trial evaluation [51]. For each trial, a candidate set of hyperparameters was sampled, the model was trained using the corresponding configuration, and the validation performance was evaluated. The objective function aimed to maximize the validation AUROC while minimizing overfitting during training epochs. The resulting optimal hyperparameter configuration was subsequently adopted for all experiments reported in this study. The hyperparameter configuration yielding the highest validation AUROC was selected as the final model for all subsequent experiments. This optimization strategy provides a systematic and reproducible approach for model selection while improving the robustness and generalization capability of HV–MultiScaleGNN across heterogeneous HV–PPIs networks.

### 4.14. Negative sampling strategy and calibration analysis

Training GNN for link prediction requires both positive and negative interactions samples to enable the model to distinguish interacting protein pairs from non-interacting ones. Since experimentally verified negative HV–PPIs are generally unavailable, negative samples were generated by randomly selecting host–virus protein pairs that were not observed in the interaction network. This uniform negative sampling strategy provides a computationally efficient and scalable approach for constructing balanced training batches while minimizing sampling bias in large heterogeneous biological networks. During each training iteration, positive host–virus interactions were combined with uniformly sampled negative protein pairs to optimize the relation-specific interaction predictor. Exposure to both interacting and non-interacting protein pairs enables HV–MultiScaleGNN to learn discriminative node representations that effectively separate true biological interactions from unlikely associations. The generated negative samples were used exclusively during training and did not overlap with the experimentally validated positive interactions.

In addition to conventional discrimination metrics, the trustworthiness of the predicted interaction probabilities was further examined using probability calibration analysis. Although metrics such as AUROC, PR-AUC, Accuracy, and F1-score measure the model’s ability to distinguish between positive and negative interactions, they do not indicate whether the predicted probabilities accurately reflect the true likelihood of an interaction. Therefore, calibration performance was assessed using the Brier Score [52] and the Expected Calibration Error (ECE) [53]. The Brier Score measures the mean squared difference between the predicted probabilities and the corresponding ground-truth labels, with lower values reflecting more accurate probabilistic predictions. Likewise, the ECE quantifies the discrepancy between prediction confidence and empirical accuracy across different confidence intervals, where smaller values correspond to better-calibrated predictions.

### 4.15. Evaluation protocol and robustness analysis

To thoroughly asses the predictive performance of HV–MultiScaleGNN, the predicted interaction logits are transformed into interaction probabilities using the sigmoid activation function during inference. These probability estimates were then employed to calculate a range of evaluation metrics, including Area Under the Receiver Operating Characteristic Curve (AUROC), Area Under the Precision–Recall Curve (AUPRC), Accuracy, Precision, Recall, F1-score, Brier Score, and ECE. Collectively, these metrics offer a comprehensive evaluation of both the discriminative capability and probability calibration of the proposed framework for HV–PPIs prediction.

The performance of HV–MultiScaleGNN was assessed under the transductive learning paradigm, in which the heterogeneous host–virus interaction graph remained unchanged across training, validation, and testing, while only the target HV–PPIs edges were split into the respective data splits. The complete heterogeneous graph consisted of 1,079,699 PPI edges, including 1,073,474 for training, 3,112 for validation, and 3,112 for testing. Overall, the network comprised 24,499 unique human proteins and 1,165 unique viral proteins, all of which were retained throughout graph representation learning. Under this evaluation strategy, only the target host–virus interaction edges were separated into different splits (the training, validation, and test sets), whereas the entire graph topology, together with all protein nodes, remained intact during representation learning. Consequently, the model was prevented from accessing unseen target interactions during training while still leveraging the full structural context of the heterogeneous interaction network.

To systematically examine the effectiveness of the proposed framework, experiments were performed using three graph representation settings: (i) Local view representation learning, which captures first-order neighborhood information; (ii) Global view representation learning, which models long-range structural dependencies through RWR and hierarchical attention; and (iii) Local+Global view, which combines both local and global representations through feature fusion. Evaluating these three settings independently enables a detailed analysis of the individual and complementary contributions of each representation strategy to HV–PPIs interaction prediction.

The evaluation protocol was further designed to examine model performance from multiple perspectives, encompassing discrimination capability, classification performance, and probability calibration. Together, these complementary evaluation criteria facilitate a rigorous comparison of the Local view, Global view, and Local+Global view representation learning strategies under identical experimental conditions. To enhance the reproducibility and reliability of the experimental findings, all models were trained using identical data partitions, optimization settings, and random initialization protocols. The same experimental configuration was maintained across all comparative studies to guarantee a fair performance comparison among the Local view, Global view, and Local+Global learning strategies. Additionally, fixed random seeds were used throughout the study to reduce variability introduced by stochastic optimization, thereby improving the consistency of the reported results.

### 4.16. Structural assessment of selected host–virus protein interactions

A downstream protein–protein docking and interface analysis was performed from the predicted three HV–PPIs pairs to investigate the structural plausibility of selected HV–PPIs identified by HV–MultiScaleGNN. One pair for each of the Local view (i.e. CYB5R3– Non-structural protein 7_SARS-CoV-2 was selected (probability = 0.8753)), Global view (i.e., EXOSC8– Non-structural protein 8_SARS-CoV-2 was selected (probability = 0.9508)). As EXOSC8–NSP8 was also the highest-ranked candidate in the Local+Global view, the second-highest-ranked non-redundant candidate, LARP7– Non-structural protein 8_SARS-CoV-2 (probability = 0.8176), was selected for this method. Thus, three distinct host–virus pairs were examined to represent the predictions obtained from the three graph representation strategies (**Table 5**). These selection criteria enabled the structural examination of candidates prioritized by different representation strategies. The host protein structures CYB5R3 (PDB ID: 7THG, chain A), EXOSC8 (PDB ID: 2NN6, chain C), LARP7 (PDB ID: 6D12, chains A and B) and virus proteins Non-structural protein 7_SARS-CoV-2 (NSP7) and Non-structural protein 8_SARS-CoV-2 (NSP8) were obtained from the NSP7–NSP8 structural complex (PDB ID: 7JLT) were obtained from the Protein Data Bank (PDB). NSP8 was represented by chain B, whereas NSP7 was represented by chains A and C, as specified for the input structures. Before docking, the coordinate files were prepared by editing the PDB files to retain the selected protein chains and remove unwanted chains and heteroatom records. The prepared structures were checked to confirm the identity of the retained protein chains and the preservation of the intended protein sequences and used as inputs for protein–protein docking. Protein–protein docking was performed using the ClusPro web server to investigate the structural compatibility of the selected host–virus PPI prediction pairs [54]. Host proteins were designated as receptors, and viral proteins were used as ligands. ClusPro employs the PIPER docking program, which performs rigid-body protein docking using a fast Fourier transform (FFT)-based search, followed by clustering of the generated docking poses to identify representative candidate complexes. Four scoring schemes, Balanced, Electrostatic-favored, Hydrophobic-favored, and van der Waals plus electrostatic (VdW+Elec) were examined. Candidate complexes were evaluated primarily according to cluster ranking and population, with the representative cluster-center models considered for further structural assessment. The predicted complexes were subsequently examined for structural plausibility and interface characteristics rather than being selected solely on the basis of individual docking scores. The selected complexes were analyzed using PDBsum1, a standalone implementation of PDBsum to characterize the predicted host–virus interfaces [55]. Hydrogen bonds, salt bridges, and non-bonded contacts were examined, and the corresponding residue identifiers, chain assignments, and reported interatomic distances were recorded. These structural features were interpreted alongside the HV–PPIs probabilities predicted by HV–MultiScaleGNN to assess the structural plausibility of the selected associations, rather than as independent experimental validation of the selected HV–PPIs. The docking and interface analyses were treated as computational assessments rather than experimental validation of biological interactions or measurements of binding affinity.

### 4.17. Gene Ontology and KEGG pathway enrichment analysis of selected host proteins

Gene Ontology (GO) enrichment and KEGG (Kyoto Encyclopedia of Genes and Genomes) pathway analyses were performed using SRplot (https://www.bioinformatics.com.cn/en), which integrates the clusterProfiler and pathview packages for functional enrichment and pathway visualization [56]. The analysis was conducted using the selected host proteins listed in **Table 5**. GO enrichment was examined across three categories: biological process (BP), cellular component (CC), and molecular function (MF) [57]. KEGG pathway enrichment was performed to identify biological pathways associated with the selected host proteins [58]. Enrichment was evaluated using the gene ratio and rich factor, as reported by the analysis platform, to describe the representation of input proteins in individual GO terms and pathways. Statistical significance was assessed using the *p*-value, with multiple-testing-adjusted *p*-values used to identify significantly enriched terms and pathways [59, 60]. An adjusted *p*-value < 0.05 was considered statistically significant.

## 5. Results

This section evaluates the predictive performance of the proposed HV– MultiScaleGNN framework for HV–PPIs prediction. First, the experimental setup and baseline methods are presented, followed by an evaluation of the three graph representation strategies. The proposed framework is then compared with representative graph representation learning methods through comprehensive performance, and hyperparameter sensitivity analyses. Finally, a case study is presented to demonstrate the biological relevance of the predicted HV–PPIs.

### 5.1. Experimental setup

All experiments were conducted in a system with NVIDIA RTX A6000 GPUs (49 GB memory). The HV–MultiScaleGNN framework was implemented using the PyTorch DL framework together with the Deep Graph Library (DGL) for efficient heterogeneous graph representation learning [61, 62]. All experiments were conducted under the CUDA v12.1 computing environment to accelerate large-scale graph processing and model training. Model optimization was carried out using the AdamW optimizer with an initial learning rate of 0.00109 and a weight decay coefficient of 1.19 × 10□□. The network parameters were optimized by minimizing the Binary Cross-Entropy with Logits (BCEWithLogitsLoss) objective function, which is well suited for binary HV–PPIs prediction. The constructed heterogeneous HV–PPIs dataset was randomly split into training, validation, and test sets at 8:1:1 ratio. To ensure reproducibility, all experiments were conducted using a fixed random seed of 42. The model was trained for up to 50 epochs with a mini-batch size of 1024. Model selection was performed on the validation set, and final predictive performance was evaluated on the independent test set. The architectural hyperparameters, including the embedding dimension, number of attention heads, dropout rate, hidden feature dimension, RWR parameters, and neighborhood sampling size, were selected through Optuna-based hyperparameter optimization. The final hyperparameter configuration corresponded to the trial achieving the highest validation AUROC and is summarized in **Table 2**

**Table 2.**
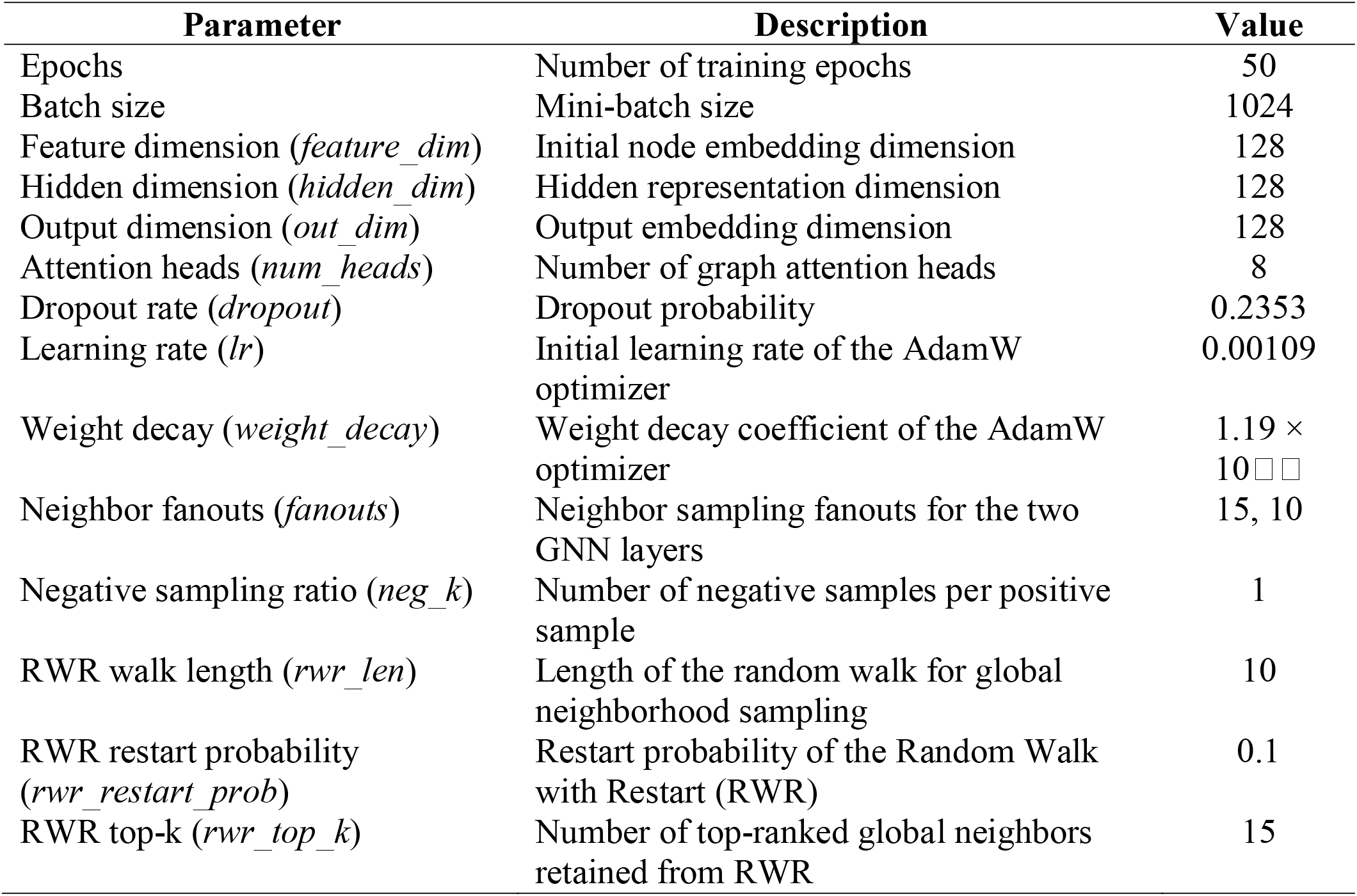
Optimal hyperparameter configuration of the proposed HV-MultiScaleGNN framework.

| Parameter | Description | Value |
|---|---|---|
| Epochs | Number of training epochs | 50 |
| Batch size | Mini-batch size | 1024 |
| Feature dimension (*feature_dim*) | Initial node embedding dimension | 128 |
| Hidden dimension (*hidden_dim*) | Hidden representation dimension | 128 |
| Output dimension (*out_dim*) | Output embedding dimension | 128 |
| Attention heads (*num_heads*) | Number of graph attention heads | 8 |
| Dropout rate (*dropout*) | Dropout probability | 0.2353 |
| Learning rate (*lr*) | Initial learning rate of the AdamW optimizer | 0.00109 |
| Weight decay (*weight_decay*) | Weight decay coefficient of the AdamW optimizer | 1.19 × 10[illegible] |
| Neighbor fanouts (*fanouts*) | Neighbor sampling fanouts for the two GNN layers | 15, 10 |
| Negative sampling ratio (*neg_k*) | Number of negative samples per positive sample | 1 |
| RWR walk length (*rwr_len*) | Length of the random walk for global neighborhood sampling | 10 |
| RWR restart probability (*rwr_restart_prob*) | Restart probability of the Random Walk with Restart (RWR) | 0.1 |
| RWR top-k (*rwr_top_k*) | Number of top-ranked global neighbors retained from RWR | 15 |

#### 5.1.1. Baseline methods

The predictive performance of HV–MultiScaleGNN framework has been compared with several widely used graph representation learning methods for HV–PPIs prediction. The selected baseline methods encompass different graph learning paradigms, including network embedding methods, homogeneous GNN, and heterogeneous GNN, thereby enabling a comprehensive evaluation of the proposed framework across diverse graph representation strategies. Specifically, the proposed framework was compared with DeepWalk [63], Node2Vec [64], Graph Convolutional Network (GCN) [32], Heterogeneous Attention Network (HAN) [47], and Heterogeneous Graph Transformer (HGT) [36]. DeepWalk and Node2Vec were included as representative network embedding methods, GCN represents convolution-based GNN, whereas HAN and HGT are representative heterogeneous GNN architectures capable of modeling multiple node and relation types. These methods provide strong and widely adopted baselines for evaluating heterogeneous biological network representation learning. For all comparative experiments, the baseline methods were implemented using their official source code or reproduced according to the configurations described in their original publications. To ensure a fair comparison, all methods were trained and tested under identical experimental conditions using the same training, validation, and testing partitions, evaluation metrics, and preprocessing pipeline. Hyperparameters for each baseline were adopted from the corresponding original publications wherever available. Otherwise, hyperparameters were tuned using the validation set within the recommended ranges to ensure a fair comparison.

### 5.2. Hyperparameter sensitivity analysis

Bayesian hyperparameter optimization was performed using the Optuna optimization framework to identify an optimal hyperparameter configuration for HV–MultiScaleGNN [51]. The optimization explored the learning rate, hidden feature dimension, output embedding dimension, dropout rate, neighborhood sampling size, number of attention heads, RWR neighborhood size, and weight decay. The objective function was defined as the mean validation AUROC obtained using five-fold cross-validation. To reduce computational cost, each fold was trained for a maximum of 50 epochs during optimization, and the average validation AUROC across the five folds guided the Bayesian search.

A total of 30 optimization trials were performed, providing a practical balance between computational efficiency and optimization effectiveness. The optimization rapidly converged to a stable high-performing region, and the best hyperparameter configuration identified during the search was adopted for all subsequent experiments. Detailed optimization analyses, including the optimization history, hyperparameter importance, parallel coordinate visualization, slice plots, and contour analyses, are provided in **Supplementary figures (Fig S1–S5)**. These analyses consistently indicate that the learning rate is the dominant factor influencing model performance, whereas the remaining architectural hyperparameters exhibit comparatively limited influence within the explored search space.

To further evaluate the robustness of the optimized solution, the distributions of the learning rate and dropout rate among the ten best-performing optimization trials are presented in **Fig. 5**. The learning rate was selected because it exhibited the highest importance during Bayesian optimization, whereas dropout represents the principal regularization parameter controlling model generalization. As illustrated in **Fig. 5 (a)**, the highest-performing trials consistently converged to learning rates near 10□^3^, despite exploring a broad hyperparameter search space. Similarly, **Fig. 5 (b)** shows that the optimal dropout values are concentrated between 0.18 and 0.24, indicating that moderate regularization provides an effective balance between minimizing overfitting and preserving discriminative feature representations.

**Fig. 5.**
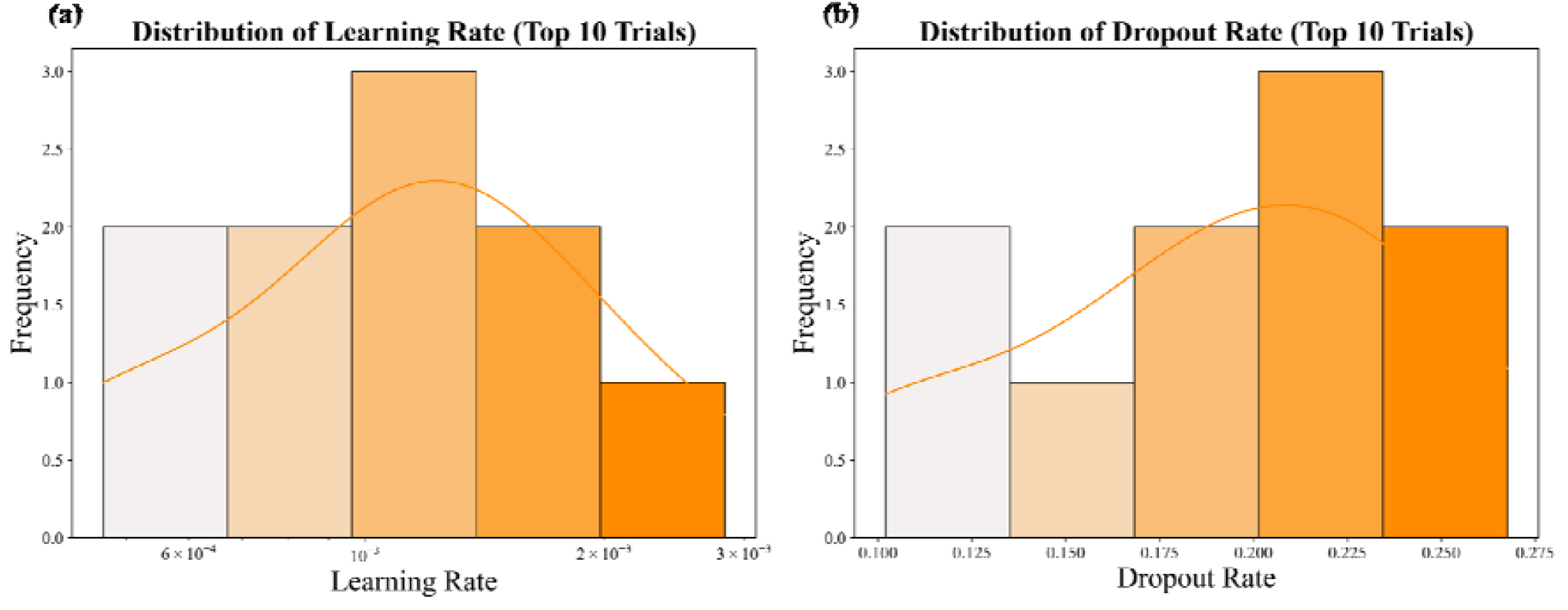
Distribution of the learning rate and dropout values among the ten best-performing Bayesian optimization trials. (a) Learning rate distribution, (b) Dropout rate distribution. The concentration of optimal configurations within relatively narrow parameter ranges indicates consistent convergence of Bayesian optimization toward reproducible high-performing solutions.

The relatively compact distributions of both hyperparameters indicate that multiple independent optimization trials converged to similar high-performing configurations rather than isolated optima. Together with the analyses presented in **Supplementary figures (Fig. S1–S5)**, these results demonstrate that HV–MultiScaleGNN exhibits stable optimization behavior and reproducible performance across the explored hyperparameter space, reducing the need for extensive hyperparameter tuning.

### 5.3. Training, validation, and test performance

To assess the generalization capability of the proposed HV–MultiScaleGNN framework, the predictive performance of the three graph representation strategies was systematically evaluated on the training, validation, and independent test datasets using the best parameters. The quantitative results are presented in **Table 3**. Since the same data partition and identical hyperparameter configuration were adopted for all representation strategies, the obtained results offer a comprehensive assessment of the learning behaviour and generalization ability of the proposed framework prior to comparison with existing methods.

**Table 3.** Predictive performance of the three graph representation strategies across the training, validation, and independent test datasets.

| **Representation** | **Dataset** | **AUROC** | **AUPRC** | **Accuracy** | **AP@20** |
|---|---|---|---|---|---|
| Global View | Train | 0.7518 | 0.7337 | 0.6822 | 0.9824 |
|  | Validation | 0.9043 | 0.8991 | 0.8451 | 0.9601 |
|  | Test | 0.9093 | 0.9147 | 0.8476 | 0.9870 |
| Local View | Train | 0.8998 | 0.8810 | 0.8196 | 1.0000 |
|  | Validation | 0.9411 | 0.9359 | 0.8707 | 1.0000 |
|  | Test | 0.9444 | 0.9422 | 0.8731 | 1.0000 |
| Global + Local View | Train | 0.8881 | 0.8667 | 0.8079 | 1.0000 |
|  | Validation | 0.9330 | 0.9296 | 0.8567 | 1.0000 |
|  | Test | 0.9378 | 0.9356 | 0.8633 | 1.0000 |

As shown in **Table 3**, all three representation strategies demonstrated consistent predictive performance across the training, validation, and independent test datasets, indicating stable learning optimization and strong generalization performance. Although the Global view yielding relatively lower performance on the training set, its validation and test results improved substantially, indicating that higher-order structural information captured via RWR effectively generalizes to previously unseen interactions. In contrast, the Local view consistently achieved the superior predictive performance across all three datasets, highlighting the strong discriminative capability of relation-aware neighborhood aggregation. The integrated Local+Global view also maintained high, consistent performance, confirming that the adaptive fusion of local and global structural information produces robust multi-scale protein representations.

Moreover, only small differences are observed between the validation and independent test performances for all three representation strategies. Such consistency suggests that the selected hyperparameter configuration and optimization strategy effectively promote good generalization while avoiding substantial overfitting. Overall, these findings indicate that the proposed HV–MultiScaleGNN framework learns reliable and transferable graph representations, providing a solid foundation for subsequent comparisons with state-of-the-art graph representation learning methods.

### 5.4. Comparison with baseline methods

All comparative experiments (DeepWalk, Node2Vec, GCN, HAN, and HGT) were conducted using the same datasets, preprocessing pipeline, and evaluation protocol to ensure a fair and unbiased comparison. The corresponding quantitative results are presented in **Table 4**. The comparative predictive performance of the proposed HV–MultiScaleGNN framework was systematically evaluated using a diverse set of complementary evaluation metrics, including the Area Under the Receiver Operating Characteristic Curve (AUROC), Area Under the Precision–Recall Curve (AUPRC), Accuracy, Average Precision at the top 20 ranked predictions (AP@20), Brier Score [52], ECE [53], Mean Reciprocal Rank (MRR), and Hits@K (Hits@1, Hits@3, and Hits@10). Together, these evaluation metrics collectively assess the discriminative capability, probability calibration, and ranking performance of the proposed framework, providing a comprehensive evaluation of HV–PPIs prediction.

**Table 4.** Comparison with state-of-the-art methods on the independent test set.

| **Model** | **AUROC** | **AUPRC** | **Accuracy** | **AP@20** | **Brier** | **ECE** | **MRR** | **Hits@1** | **Hits@3** | **Hits@10** |
|---|---|---|---|---|---|---|---|---|---|---|
| DeepWalk | 0.9184 | 0.9092 | 0.8497 | 1.0000 | 0.1137 | 0.0422 | 0.9536 | 0.9072 | 1.0000 | 1.0000 |
| Node2Vec | 0.9186 | 0.9121 | 0.8477 | 1.0000 | 0.1136 | 0.0372 | 0.9528 | 0.9056 | 1.0000 | 1.0000 |
| HAN | 0.9250 | 0.9168 | 0.8553 | 1.0000 | 0.1076 | 0.0354 | 0.9573 | 0.9146 | 1.0000 | 1.0000 |
| HGT | 0.9038 | 0.9050 | 0.8204 | 1.0000 | 0.1381 | 0.0818 | 0.9484 | 0.8969 | 1.0000 | 1.0000 |
| GCN | 0.9285 | 0.9253 | 0.8583 | 1.0000 | 0.1055 | 0.0241 | 0.9555 | 0.9110 | 1.0000 | 1.0000 |
| **Local view** | **0.9444** | **0.9422** | **0.8731** | **1.0000** | **0.1005** | **0.0659** | **0.9659** | **0.9319** | **1.0000** | **1.0000** |
| **Glocal view** | **0.9093** | **0.9147** | **0.8476** | **0.9870** | **0.1156** | **0.0379** | **0.9547** | **0.9094** | **1.0000** | **1.0000** |
| **Local+Global view** | **0.9378** | **0.9356** | **0.8633** | **1.0000** | **0.1034** | **0.0558** | **0.9624** | **0.9248** | **1.0000** | **1.0000** |

The proposed HV–MultiScaleGNN framework exhibited robust predictive performance across its three graph representation strategies: the Local view, Global view, and the integrated Local+Global view. Although each strategy captures distinct structural characteristics of the heterogeneous interaction network, all three achieved strong predictive performance on the independent test set. Among these approaches, the Local view obtained the best overall results, achieving the highest AUROC (0.9444), AUPRC (0.9422), Accuracy (0.8731), and MRR (0.9659). These findings suggest that information derived from immediate neighborhood interactions provides highly informative and discriminative representations for HV–PPIs prediction. In comparison, the Global view also produced competitive results (AUROC = 0.9093 and AUPRC = 0.9147), indicating that higher-order structural dependencies captured through RWR preserve biologically relevant information even in the absence of direct neighborhood aggregation. The integrated Local+Global view demonstrated consistently balanced performance across all evaluation metrics (AUROC = 0.9378, AUPRC = 0.9356, Accuracy = 0.8633), suggesting that the integration of local and global topological information generates comprehensive multi-scale protein representation with strong generalization capability across heterogeneous biological networks.

A comparison with other graph representation learning methods further highlights the effectiveness of the proposed framework. The Local view consistently outperformed all baseline methods across AUROC, AUPRC, Accuracy, and MRR (**See Table 4**). In addition to conventional classification metrics, calibration- and ranking-oriented evaluations provide further evidence of the robustness of the proposed framework. The Global view exhibited the lowest ECE = 0.0379, indicating that it generated the most reliable probability estimates among the three representation strategies. Conversely, the Local view achieved the lowest Brier Score (0.1005) and the highest Mean Reciprocal Rank (MRR = 0.9659), demonstrating superior probabilistic accuracy and excellent ranking performance. The integrated Local+Global view produced intermediate calibration results (Brier Score = 0.1034 and ECE = 0.0558) while consistently maintaining strong ranking capability (MRR = 0.9624). Furthermore, all three representation strategies achieved perfect scores for Hits@3 and Hits@10, indicating that experimentally validated HV–PPIs were consistently ranked among the highest-confidence predictions generated by the framework. Overall, these results highlight the flexibility, robustness, and practical applicability of the proposed framework for large-scale HV–PPIs prediction and candidate interaction prioritization.

### 5.5. Case study

To further evaluate the biological relevance, robustness, and generalization capability of HV–MultiScaleGNN, an independent external validation study was conducted using 67 experimentally validated HV–PPIs collected from the published literature. The complete external validation dataset is provided in **Supplementary file S3**. The interaction probability for each candidate HV–PPIs was computed as

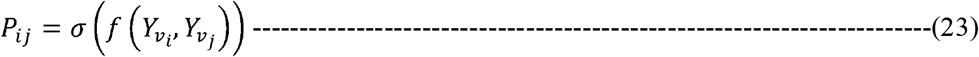

where 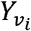 and 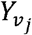 denote final protein embeddings of the host and viral proteins generated by the selected graph representation strategy (Local view, Global view, or Local+Global view), *f*(·) represents the relation-specific MLP interaction decoder, and *σ* (·) denotes the sigmoid activation function that converts the decoder output into an interaction probability ranging from 0 to 1. For the Local+Global view, these correspond to the unified protein embeddings obtained through multi-scale feature fusion, whereas for the Local view and Global view, they correspond to the respective branch-specific protein embeddings. Larger values of *P*_*ij*_ indicate greater confidence that the corresponding HV–PPIs pair forms a biologically plausible interaction.

For biological interpretation, only high-confidence predictions with a prediction probability greater than or equal to 0.70 were considered. Representative high-confidence experimentally validated HV–PPIs recovered by the proposed framework are summarized in **Table 5**, whereas the complete prediction results, including the predicted interaction probabilities for all 67 experimentally validated external HV–PPIs, are provided in **Supplementary file S4**. Rather than representing competing prediction models, the three graph representation strategies capture complementary structural characteristics of heterogeneous biological networks. Several host proteins were assigned consistently high prediction probabilities for interactions with biologically important viral proteins, including Non-structural protein 7_SARS-CoV-2, Non-structural protein 8_SARS-CoV-2, Membrane protein_SARS-CoV-2, ORF10 protein_SARS-CoV-2, and Glycoprotein G_Nipah virus. These viral proteins have well-established roles in viral replication, assembly, host interaction, and cell entry [65]. Non-structural proteins 7 and 8 of SARS-CoV-2 function as essential cofactors of the viral RNA-dependent RNA polymerase complex and are indispensable for viral RNA synthesis and genome replication [85]. Membrane protein_SARS-CoV-2 is the principal structural protein responsible for virion assembly, morphogenesis, and viral particle formation [66]. ORF10 protein_SARS-CoV-2 has been implicated in host ubiquitination pathways and modulation of host cellular processes [87], whereas Glycoprotein G_Nipah virus mediates viral attachment to ephrin-B2 and ephrin-B3 receptors and is a major determinant of viral entry and host tropism [67].

**Table 5.** Representative high-confidence experimentally validated HV–PPIs recovered by the proposed framework.

| Host protein (Gene ID) | Viral protein | Local view | Global view | Local+Global view |
|---|---|---|---|---|
| EXOSC8 (11340) | Non-structural protein 8_SARS-CoV-2 | **0.7908** | **0.9508** | **0.8251** |
| LARP7 (51574) | Non-structural protein 8_SARS-CoV-2 | **0.7953** | **0.9156** | **0.8176** |
| MEPCE (56257) | Non-structural protein 8_SARS-CoV-2 | **0.8004** | **0.8742** | **0.8047** |
| NSD2 (7468) | Non-structural protein 8_SARS-CoV-2 | **0.7577** | **0.9197** | **0.7894** |
| MRPS5 (64969) | Non-structural protein 8_SARS-CoV-2 | **0.7625** | **0.9185** | **0.7845** |
| CCDC86 (79080) | Non-structural protein 8_SARS-CoV-2 | **0.7725** | **0.9147** | **0.7307** |
| CYB5B (80777) | Non-structural protein 7_SARS-CoV-2 | **0.8520** | 0.5635 | **0.7924** |
| RAB2A (5862) | Non-structural protein 7_SARS-CoV-2 | **0.7985** | 0.6044 | **0.7863** |
| FAM162A (26355) | Non-structural protein 7_SARS-CoV-2 | **0.7827** | 0.5797 | **0.7696** |
| CYB5R3 (1727) | Non-structural protein 7_SARS-CoV-2 | **0.8753** | 0.4935 | **0.7041** |
| BZW2 (28969) | Membrane protein_SARS-CoV-2 | **0.7655** | 0.3248 | **0.7309** |
| REEP5 (7905) | Membrane protein_SARS-CoV-2 | **0.7725** | 0.6170 | **0.7104** |
| EFNB2 (1948) | Glycoprotein G_Nipah virus | **0.8037** | 0.2624 | 0.5251 |
| ELOC (6921) | ORF10 protein_SARS-CoV-2 | **0.7725** | 0.2067 | 0.6631 |
| RBX1 (9978) | ORF10 protein_SARS-CoV-2 | **0.7577** | 0.2442 | 0.6607 |

\*The probability values ≥ 0.70 are highlighted in bold.

Collectively, these results demonstrate that HV–MultiScaleGNN can successfully recover biologically meaningful HV–PPIs that were absent from the training graph, supporting its potential as a topology-driven framework for prioritizing novel host–virus interactions for future experimental validation.

### 5.6. Attention weight analysis and interpretability

To uncover the relationships between specific viral proteins and their corresponding human proteins, we analyzed the attention coefficients learned by the Local view model, that achieved the best predictive performance among the three graph representation strategies. The corresponding attention visualizations for selected representative interactions are presented in **Supplementary figures (Fig. S6)**. Importantly, several highly attended neighboring proteins identified by the Local view model are supported by independent biological evidence, demonstrating the biological relevance of the learned attention patterns. For example, ACSL3 (NCBI Gene ID: 2181), which consistently received the highest attention weights for Non-structural protein 7_SARS-CoV-2 associated interactions, has been experimentally implicated in coronavirus lipid metabolism and viral replication [68]. Likewise, MAP7D1 (NCBI Gene ID: 55700), repeatedly prioritized for ORF10 protein_SARS-CoV-2, has been reported as a human interaction partner of ORF10 protein_SARS-CoV-2 [69]. Similarly, SRP72 (NCBI Gene ID: 6731), one of the most highly attended proteins associated with Non-structural protein 8_SARS-CoV-2, has been experimentally identified as an NSP8-interacting host factor involved in the SRP/Sec61-mediated protein translocation pathway [84]. Together, these observations indicate that the Local view model captures biologically meaningful local topological patterns rather than assigning attention arbitrarily, thereby providing an interpretable explanation for its superior performance in host–virus protein–protein interaction prediction.

### 5.7. ClusPro-based structural assessment and interfacial interactions of selected host– virus protein complexes

The three HV–PPIs pairs prioritized by HV–MultiScaleGNN were further examined using ClusPro to assess the structural plausibility of their predicted associations. Candidate docking models were critically evaluated using the Balanced, Electrostatic-favored, Hydrophobic-favored, and van der Waals plus electrostatic (VdW+Elec) scoring schemes. The top-ranked candidate models were examined in PyMOL to assess the relative orientation of the interacting proteins and the geometry of their predicted interfaces. Models with distinct interface characteristics were also evaluated using PDBsum1 to characterize residue-level contacts, including hydrogen bonds, salt bridges, and non-bonded interactions. The final models were selected by considering cluster information and the overall structural plausibility of the predicted complexes rather than relying exclusively on individual docking scores or contact counts. The resulting three-dimensional complex structures and their corresponding interface visualizations are presented in **Fig. 6 (a), (b) and (c)**, respectively.

**Fig. 6.**
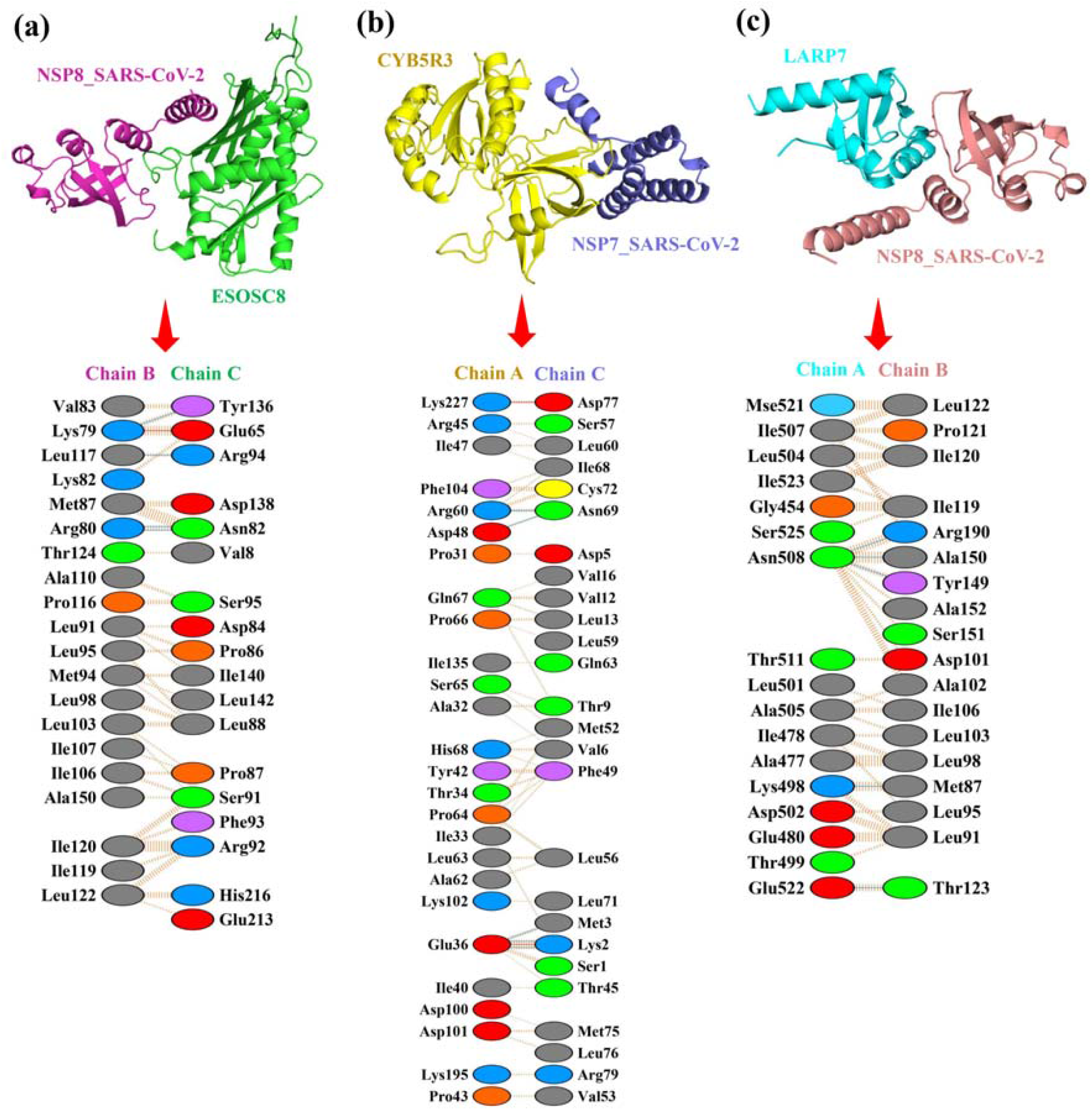
Structural models and residue-level interfacial interactions of three selected HV– PPIs complexes predicted by HV–MultiScaleGNN. The upper panels show the three-dimensional representations of the selected docked complexes, while the corresponding lower panels illustrate residue-level interactions at the predicted interfaces obtained using PDBsum1. (a) EXOSC8 (chain B)–NSP8_SARS-CoV-2, (b) CYB5R3 (chain A)–SARS-CoV-2 NSP7 (chain C), and (c) LARP7 (chain A)–SARS-CoV-2 NSP8 (chain B). The lower interaction diagrams display the participating amino acid residues and their interfacial contacts. These structural representations provide a visualization of the predicted docking poses and residue-level interaction patterns, including hydrogen bonds, salt bridges, and non-bonded contacts, as identified by the interface analysis.

PDBsum1 analysis of the selected EXOSC8–NSP8 model identified four hydrogen bonds, one salt bridge, and 90 non-bonded contacts across the predicted interface. The reported hydrogen bonds involved Arg80 of EXOSC8 and Asn82 of NSP8, with two contacts at distances of 2.72 and 3.29 Å; Leu117 of EXOSC8 and Arg94 of NSP8, at 3.06 Å; and Lys79 of EXOSC8 and Tyr136 of NSP8, at 2.50 Å. The analysis also identified a salt bridge between Lys79 of EXOSC8 and Glu65 of NSP8, with a reported distance of 2.47 Å. The recurrence of contacts involving Lys79 of EXOSC8 and Glu65 of NSP8, together with the hydrogen-bonding interactions involving Arg80 and Asn82, highlights several polar and electrostatic features of the modeled interface. In addition, the 90 reported non-bonded contacts indicate an extensive set of close interatomic contacts within the selected pose. Collectively, these observations define a predicted interface involving multiple polar interactions and a salt bridge, providing residue-level hypotheses for further investigation of the EXOSC8–NSP8 association.

The selected CYB5R3–NSP7 model contained five hydrogen bonds, two salt bridges, and 128 non-bonded contacts, representing the highest reported numbers of both salt bridges and non-bonded contacts among the three selected complexes. The hydrogen-bonding interactions involved Glu36 of CYB5R3 and Lys2 of NSP7, with reported distances of 2.65 and 2.45 Å; Glu36 of CYB5R3 and Met3 of NSP7, at 2.89 Å; Asp48 of CYB5R3 and Asn69 of NSP7, at 3.12 Å; and Arg60 of CYB5R3 and Asn69 of NSP7, at 2.63 Å. The two reported salt bridges connected Glu36 of CYB5R3 with Lys2 of NSP7, at 2.45 Å, and Lys227 of CYB5R3 with Asp77 of NSP7, at 2.77 Å. These interactions identify two potential electrostatic contact regions, accompanied by hydrogen bonds involving Glu36, Asp48, Arg60, Lys2, Met3, and Asn69. The comparatively large number of non-bonded contacts further describes the close packing observed in this particular docked configuration. Together, the predicted hydrogen bonds and salt bridges provide a detailed description of the selected CYB5R3–NSP7 interface and identify residues that may be useful for subsequent structural investigation.

The selected LARP7–NSP8 model contained five hydrogen bonds and 76 non-bonded contacts; PDBsum1 did not report a salt bridge for this complex. The hydrogen bonds involved Lys498 of LARP7 and Met87 of NSP8, at a distance of 2.76 Å; Asn508 of LARP7 and Arg190 of NSP8, at 2.69 Å; Asn508 of LARP7 and Tyr149 of NSP8, at 2.92 Å; Asn508 of LARP7 and Ala150 of NSP8, at 2.71 Å; and Glu522 of LARP7 and Thr123 of NSP8, at 3.01 Å. Asn508 of LARP7 participated in three of the five reported hydrogen bonds, making it a notable residue within the predicted interface. The contacts involving Lys498, Asn508, and Glu522 of LARP7, together with Met87, Arg190, Tyr149, Ala150, and Thr123 of NSP8, indicate a network of predicted polar interactions across the interface. The 76 reported non-bonded contacts provide additional information about the close interatomic packing of the selected complex. These findings characterize the modeled LARP7–NSP8 interface and identify candidate residues for further investigation.

Overall, the ClusPro-based structural assessment provided residue-level descriptions of the predicted EXOSC8–NSP8, CYB5R3–NSP7, and LARP7–NSP8 interfaces, complementing the host–virus interaction predictions generated by HV–MultiScaleGNN. The selected models contained identifiable hydrogen bonds, non-bonded contacts, and salt bridges. These features support the structural plausibility of the selected poses as computational hypotheses.

### 5.8. Functional enrichment of predicted HV–PPIs partners

GO enrichment analysis of the host proteins revealed functional associations with RNA metabolism, ribonucleoprotein complexes, ubiquitin-ligase machinery, oxidoreductase activity etc (**Table 5**). In the biological process (BP) category, snRNA metabolic process was the most significantly enriched term, with EXOSC8, LARP7, and MEPCE as the contributing proteins. All three host proteins were associated with SARS-CoV-2 non-structural protein 8 (NSP8) in the predicted interaction presented in **Table 5**. The enrichment is consistent with established roles of these proteins in RNA processing and regulation. EXOSC8 is a structural component of the RNA exosome, a conserved multiprotein complex involved in RNA processing, surveillance, and degradation, including the maturation and turnover of non-coding RNAs [70]. LARP7 and MEPCE are core components of the 7SK small nuclear ribonucleoprotein (7SK snRNP), which regulates RNA polymerase II transcription elongation through its control of positive transcription elongation factor b (P-TEFb) activity [71]. Additional enriched terms included regulation of protein localization to Cajal bodies, snRNA transcription, and non-coding RNA 3′-end processing. The involvement of LARP7 and MEPCE is consistent with evidence that the 7SK snRNP participates in the regulation of snRNA and snoRNA gene transcription, in addition to its role in controlling transcription elongation [71]. These annotations provide functional context for the EXOSC8, LARP7, and MEPCE-associated interaction predictions but do not establish that NSP8 directly influences RNA processing or transcription in infected cells. In the cellular component (CC) category, the Cullin 2–RING ubiquitin ligase complex was significantly enriched, with ELOC and RBX1 as the overlapping proteins. Both proteins were also associated with SARS-CoV-2 non-structural protein 8 (NSP8) in the predicted interaction presented in **Table 5**. ELOC encodes Elongin C, a component of the Cullin-RING E3 ubiquitin ligase machinery, whereas RBX1 is a RING-box protein that participates in ubiquitin transfer by recruiting the E2 ubiquitin-conjugating enzyme to the ligase complex. The Cullin 2–Elongin B–Elongin C– RBX1 machinery is involved in substrate ubiquitination and proteasomal degradation, including the regulation of hypoxia-inducible factor 1α (HIF-1α) through the von Hippel– Lindau protein-containing E3 ligase complex [72, 73]. The small nuclear ribonucleoprotein complex and Sm-like protein family complex were also enriched, with LARP7 and MEPCE contributing to both terms. In addition, the oxidoreductase complex was enriched through CYB5B and CYB5R3, both interaction with SARS-CoV-2 NSP7 in **Table 5**. Cytochrome b5 reductase 3 (CYB5R3) catalyses NADH-dependent electron transfer, while cytochrome b5 proteins participate in redox reactions supporting several cellular enzymatic processes [74]. Together, these findings indicate that the selected host-protein set encompasses several distinct functional groups rather than a single dominant cellular process. For molecular function (MF), snRNA binding was significantly enriched, with LARP7 and MEPCE as the contributing proteins. This result is consistent with their established roles in the 7SK snRNP, in which LARP7 binds 7SK RNA and cooperates with MEPCE to maintain the stability and integrity of the ribonucleoprotein complex [71]. S-adenosylmethionine-dependent methyltransferase activity was also enriched at the nominal level, with MEPCE and NSD2 contributing to the term. MEPCE is a methylphosphate capping enzyme involved in 7SK RNA metabolism, whereas NSD2 is a histone methyltransferase associated with chromatin regulation. CYB5R3 contributed to the oxidoreductase-related molecular-function annotation, consistent with its established role in cellular electron transfer [74]. These functional annotations support the biological relevance of the selected host-protein set, although the enrichment analysis alone cannot establish whether the predicted viral partners alter the activities of these proteins. The corresponding GO enrichment results for biological process (BP), cellular component (CC), and molecular function (MF) are presented in **Fig. 7 (a)**.

**Fig. 7.**
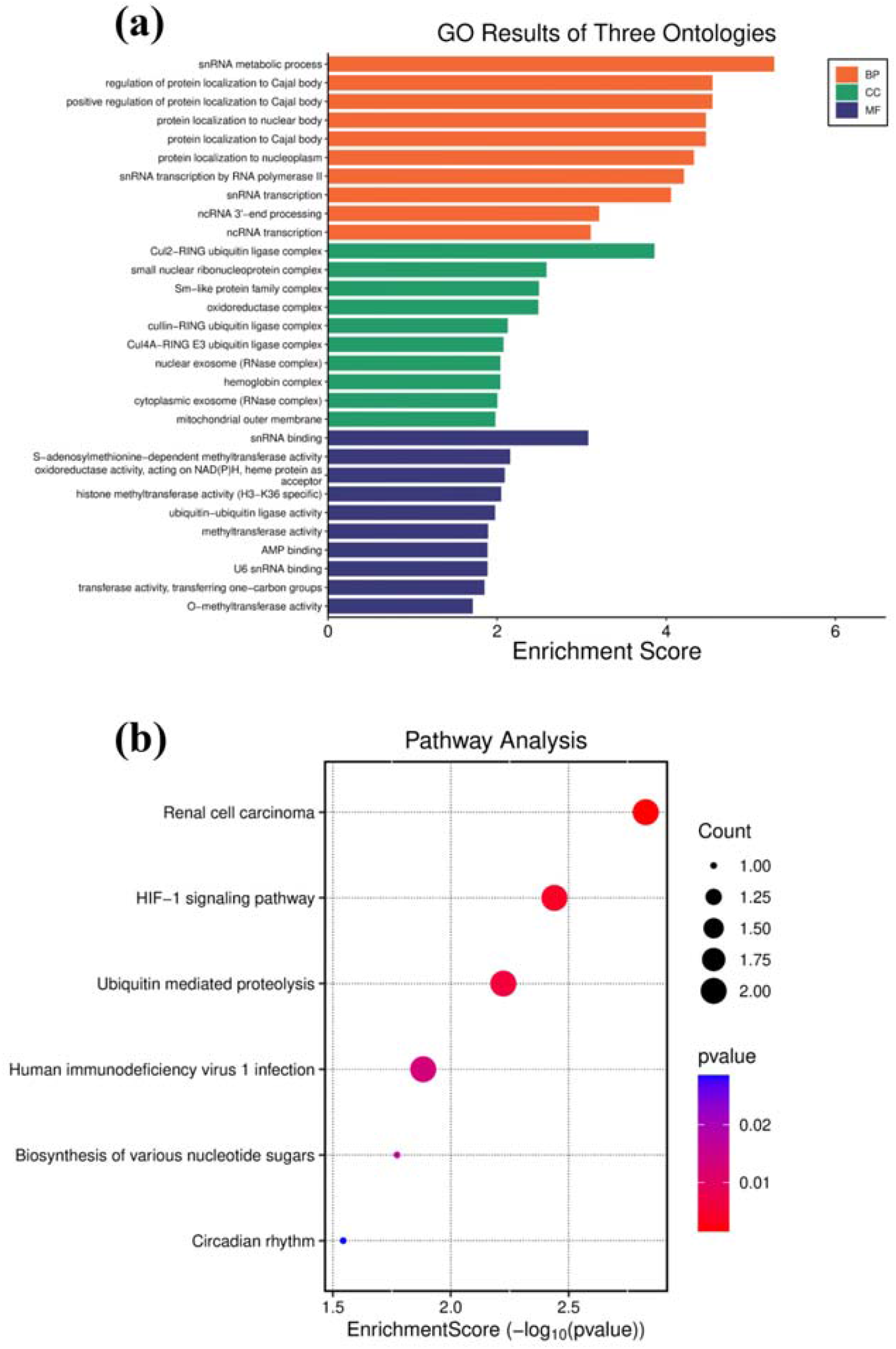
Gene Ontology (GO) and Kyoto Encyclopedia of Genes and Genomes (KEGG) pathway enrichment analysis of the selected host proteins listed in Table 5. (a) GO enrichment results across the three ontologies: biological process (BP), cellular component (CC), and molecular function (MF). The horizontal bars represent the Enrichment Scores of the enriched GO terms, (b) KEGG pathway enrichment analysis shows the enriched pathways, with the x-axis representing the Enrichment Score. The size of each bubble represents the number of input host proteins associated with the corresponding pathway, while the colour indicates the p-value. The analyses were performed using the selected host proteins prioritized by HV–MultiScaleGNN.

KEGG pathway enrichment analysis identified several pathways, including renal cell carcinoma, HIF-1 signaling pathway, ubiquitin-mediated proteolysis, human immunodeficiency virus 1 (HIV-1) infection, biosynthesis of various nucleotide sugars, and circadian rhythm (**Fig. 7 (b)**). Renal cell carcinoma, HIF-1 signaling, ubiquitin-mediated proteolysis, and HIV-1 infection shared ELOC and RBX1 as the contributing host proteins, both of which were predicted to interact with SARS-CoV-2 ORF10 (**Table 5**). The involvement of ELOC and RBX1 in these pathway annotations is biologically relevant because both proteins are components of the Cullin-RING ubiquitin ligase machinery, which participates in ubiquitin-dependent protein regulation and proteasomal degradation [96][73]. In particular, the CUL2–RBX1–Elongin B–Elongin C–VHL complex regulates the ubiquitination and proteasomal degradation of hypoxia-inducible factor 1α (HIF-1α), providing a mechanistic link between ubiquitin-mediated proteolysis and HIF-1-associated cellular regulation [73]. This functional association provides a biological context for investigating the predicted SARS-CoV-2 ORF10–ELOC and ORF10–RBX1 interactions in relation to host ubiquitin-mediated proteolysis and HIF-1-associated pathways. Interestingly, previous experimental work demonstrated that SARS-CoV-2 ORF10 interacts with the ZYG11B substrate receptor of the CUL2 ubiquitin ligase complex and enhances its E3 ligase activity, promoting ubiquitination and proteasomal degradation of the host protein IFT46 [75]. Furthermore, CYB5R3, which was predicted to interact with SARS-CoV-2 non-structural protein 7 (NSP7), contributed to the biosynthesis of various nucleotide sugars pathway, whereas RBX1 contributed to the circadian rhythm pathway. These observations indicate that the selected host proteins are associated with diverse cellular pathway annotations. Nevertheless, these enrichment findings reflect the existing functional annotations of the host proteins and do not establish direct viral regulation or functional consequences of the predicted interactions.

Collectively, the GO and KEGG enrichment analyses revealed distinct functional groupings among the host proteins prioritized by HV–MultiScaleGNN. These functional groupings are broadly consistent with the established cellular roles of the respective host proteins and provide additional biological context for the predicted HV–PPIs. Nevertheless, the enrichment findings reflect existing functional annotations of the host proteins, and not all enriched terms necessarily have a direct functional relationship with HV–PPIs. Experimental investigations will be required to establish the biological relevance and potential functional consequences of the predicted interactions.

## 6. Discussion

The present study shows that heterogeneous graph topology provides sufficient structural information to support accurate HV–PPIs prediction without relying on explicit protein sequence, structural, or phenotype-derived features. Unlike most existing computational approaches that learn representations from molecular characteristics of individual proteins, the proposed HV–MultiScaleGNN framework formulates HV–PPIs prediction as a topology-driven heterogeneous link prediction problem and systematically investigates three complementary graph representation strategies: Local view, Global view, and an integrated Local+Global view. This unified framework enables a systematic comparison of how different graph propagation scales influence prediction performance, probability calibration, and interaction ranking, providing new insights into topology-driven representation learning for heterogeneous biological networks.

The experimental results indicate that the three representation strategies capture complementary aspects of biological network organization rather than redundant structural information. The Local view achieved the highest discrimination performance, suggesting that immediate molecular neighborhoods provide highly informative signals for identifying direct host–virus interactions. This observation is biologically reasonable, as viral proteins frequently hijack specific host protein complexes and signaling pathways, making localized interaction patterns particularly informative for predicting direct molecular associations. In contrast, the Global view consistently produced the lowest ECE, suggesting that higher-order topological relationships enhance the reliability of predicted interaction probabilities by incorporating broader functional context across the interaction network. The integrated Local+Global view combined these complementary representations and achieved consistently balanced performance across discrimination, calibration, and ranking metrics, demonstrating the complementary nature of local and global graph representations. Importantly, these findings indicate that no single representation strategy is universally optimal. Instead, each captures distinct biological characteristics, highlighting the value of evaluating multiple structural scales within the unified HV–MultiScaleGNN framework.

Recent HV-PPI prediction methods have increasingly relied on DL and protein language models, including DeepViral, DeepVHPPI, DeepHVI, HBFormer, and ECA-PHV, which utilize protein sequences, structural descriptors, evolutionary information, phenotype data, and transformer-derived embeddings [27–31]. These approaches have substantially improved predictive performance by learning informative molecular representations. However, as they primarily construct representations from individual proteins, they may not fully capture the systems-wide organization of host–virus interaction networks. In contrast, HV– MultiScaleGNN learns representations directly from heterogeneous graph topology, demonstrating that network organization itself encodes substantial predictive information. Rather than replacing sequence-based or protein language model approaches, the proposed framework provides an orthogonal source of biological information by modeling functional organization within cellular interaction networks. Consequently, integrating topology-driven graph representations with protein language models and structural information may represents a promising direction for future multimodal HV–PPIs prediction.

Compared with classical network embedding methods such as DeepWalk and Node2Vec, which generate topology-preserving but task-independent node embeddings, HV– MultiScaleGNN performs end-to-end task-specific representation learning optimized directly for HV–PPIs prediction. Furthermore, although modern GNN architectures, including GCN, GAT, HAN, and HGT, have significantly advanced biological graph representation learning through neighborhood aggregation and relation-aware message passing, most existing approaches rely on a single propagation mechanism. Local message passing effectively captures immediate molecular interactions but may overlook long-range functional dependencies, whereas extensive graph propagation can introduce information dilution and over-smoothing. Instead of assuming that one propagation strategy is universally optimal, HV–MultiScaleGNN systematically explores complementary local, global, and integrated graph representations within a unified framework, yielding a more comprehensive understanding of how different propagation scales influence to biological interaction prediction.

To further examine the biological relevance of the predicted interactions, selected HV– PPIs were subjected to protein–protein docking and functional enrichment analyses. The ClusPro-based assessment identified distinct residue-level interaction patterns involving hydrogen bonds, salt bridges, and non-bonded contacts, providing structural hypotheses for the predicted associations. In parallel, GO and KEGG enrichment analyses highlighted functional annotations related to RNA-associated processes, oxidoreductase complexes, and ubiquitin-mediated proteolysis among the selected host proteins. These complementary analyses provide additional structural and functional context for the predictions generated by HV–MultiScaleGNN. Experimental validation will therefore be necessary to establish the biological relevance and potential functional consequences of these predicted associations.

The comparative analyses also highlight the practical value of evaluating prediction quality beyond conventional discrimination metrics. This comprehensive evaluation is particularly important for biological applications, where computational predictions are used to prioritize candidates for costly experimental validation. Reliable confidence estimates and accurate interaction ranking therefore complement traditional discrimination performance and increase the practical utility of computational HV–PPIs prediction. The observed differences among the three graph representation strategies further imply that HV–PPIs are organized across multiple structural scales within the cellular interaction network. The superior discrimination achieved by the Local view indicates that direct molecular neighborhoods provide highly informative signals for identifying experimentally validated host–virus interactions, whereas the improved calibration of the Global view suggests that higher-order topological relationships contribute to more reliable confidence estimation. These findings indicate that local and global graph structures encode complementary biological information and justify the systematic evaluation of multiple representation strategies within the HV– MultiScaleGNN framework rather than relying on a single propagation mechanism.

Overall, the findings suggest that heterogeneous graph topology provides a valuable and complementary source of biological information for HV–PPIs prediction. By systematically assessing three graph representation strategies within a unified computational framework, HV–MultiScaleGNN offers new insights into the respective roles of local neighborhood aggregation and higher-order network organization in heterogeneous biological networks. Rather than competing with recent sequence-based or transformer-driven methodologies, the framework complements these approaches and positions topology-driven multi-scale graph representation learning as a promising direction for future HV–PPIs prediction. As increasingly comprehensive interaction databases, protein language models, structural biology resources, and multimodal biological datasets become available, integrating these complementary information sources is expected to further improve the accuracy, robustness, and biological interpretability of next-generation host–virus interaction prediction frameworks.

## 7. Limitations and future directions

Although the proposed HV–MultiScaleGNN framework demonstrates strong predictive performance using only the topology of a heterogeneous host–virus interaction network, several limitations remain. First, the framework is evaluated under a transductive learning setting in which all proteins are available during training. Future work should investigate inductive heterogeneous GNN architecture capable of generalizing to previously unseen host or viral proteins. Second, unlike many recent HV–PPIs prediction methods that incorporate protein sequences, structural information, or protein language model embeddings, the proposed framework relies exclusively on heterogeneous network topology. While this topology-driven design demonstrates that network structure alone contains substantial predictive information, integrating complementary molecular features such as protein sequences, structural representations, and functional annotations may further improve predictive performance, particularly for sparsely connected or newly identified proteins. Third, the current heterogeneous interaction network represents a static snapshot of biological systems and therefore cannot capture the temporal and context-dependent dynamics of host– virus interactions during infection.

The predictive performance of HV–MultiScaleGNN also depends on the completeness and quality of available biological interaction databases, which remain incomplete and are often biased toward extensively studied viruses. Furthermore, the structural assessment was restricted to three selected HV–PPIs pairs. Although the docking analyses provided computationally predicted interaction poses and residue-level interface information, these findings do not establish physical interactions or experimentally measured binding affinities. Similarly, the GO and KEGG enrichment analyses were based on a limited set of selected host proteins and therefore provide preliminary functional context rather than comprehensive evidence of virus-associated pathway modulation. In addition, although the proposed attention mechanism provides an interpretable view of neighborhood importance, attention weights should not be interpreted as causal explanations of biological interactions, and computational predictions ultimately require experimental validation. Future research will therefore focus on extending the proposed topology-driven multi-scale framework through inductive and dynamic graph learning, multimodal integration with sequence and structural information, improved explainable artificial intelligence, and cross-species or pan-viral learning to further enhance its robustness, generalizability, and biological applicability. Experimental investigation of selected high-confidence predictions will also be important for establishing their biological relevance.

## 6. Conclusions

In this study, we introduced HV–MultiScaleGNN, a topology-driven multi-scale GNN framework for predicting HV–PPIs. Unlike conventional GNN approaches that rely on a single graph representation strategy, the proposed framework systematically examines the complementary roles of Local view, Global view, and integrated Local+Global view representations within a unified heterogeneous graph learning framework. Extensive experimental evaluation of the three graph representation strategies implemented in the HV– MultiScaleGNN framework revealed consistent and reliable predictive performance across multiple evaluation metrics while preserving strong generalization on independent test data. Among these strategies, the Local view achieved the highest discrimination performance, the Global view exhibited superior probability calibration. Meanwhile, the integrated Local+Global view achieved a balanced performance across predictive discrimination, interaction ranking, and calibration metrics. Collectively, these findings indicate that local and global graph representations capture unique yet complementary biological information, and their combination results in balanced improvements across predictive discrimination, interaction ranking, and probability calibration. The results further demonstrate that heterogeneous graph topology alone contains substantial predictive information and that multi-scale representation learning provides a systematic approach for modeling complex host–virus interaction networks. Complementary structural assessment of three selected host– virus pairs identified predicted residue-level interfacial interactions, while GO and KEGG enrichment analyses provided additional functional context for the selected host proteins. These downstream analyses offer structural and functional perspectives on selected model predictions but do not constitute experimental confirmation of the predicted interactions or demonstrate direct viral regulation of the enriched biological pathways. Although the current framework focuses on topology-driven representation learning within a transductive setting, it provides a flexible foundation for future developments involving inductive graph learning, protein language models, structural biology, multimodal biological data, and explainable artificial intelligence. With the continuous growth of biological interaction resources, the integration of these complementary information sources is anticipated to further improve both the predictive capability and the biological interpretability of graph-based models.

Overall, HV–MultiScaleGNN demonstrates that leveraging graph representations across multiple structural scales serves as an effective computational framework for HV–PPIs prediction. In addition to its application to HV–PPIs prediction, the proposed multi-scale graph learning paradigm represents a general strategy for modeling complex biological networks and may facilitate a wide range of biomedical applications, including drug–target interaction prediction, disease gene prioritization, network medicine, and infectious disease research.

## Acknowledgments

DBT is thanked for the financial support in the form of the Centre of Excellence in Advanced Computation and Data Sciences (Ref. No: BT/PR40188/BTIS/137/27/2021) and National Networking Project (NNP) (Ref. No: BT/PR40233/BTIS/137/74/2023). Manuscript communication no: CSIR-NEIST/PUB/2026/348.

## Data availability

The source code and data used in this study are available upon request.

## AI Disclosure

During the preparation of this work the author(s) used ChatGPT (https://chat.openai.com/) in order to improve the language quality and making it concise. The text was then reviewed and edited for inclusion in the content. After using this tool/service, the author(s) reviewed and edited the content as needed and take(s) full responsibility for the content of the publication.

