## Supplementary figure for "HV–MultiScaleGNN: A Multi-Scale Graph Neural Network Framework for Host–Virus Protein–Protein Interactions Prediction"

**[*Supplementary figures*]**

**Fig. S1**. Optimization history of Bayesian hyperparameter optimization using Optuna. Each blue point represents the validation AUROC obtained in an individual optimization trial, while the red line indicates the best objective value achieved up to each trial.

**Fig. S2**. Relative importance of optimized hyperparameters estimated by Optuna. Importance values indicate the contribution of each hyperparameter to the variation in validation AUROC across the 30 Bayesian optimization trials. The learning rate was the dominant factor influencing predictive performance, whereas the remaining hyperparameters exhibited comparatively low importance within the explored search space.

**Fig. S3. Parallel coordinate plot illustrating the hyperparameter configurations explored during Bayesian optimization. Each polyline represents one optimization trial, while the color indicates the validation AUROC (objective value), with darker lines corresponding to better-performing trials. The visualization shows that high-performing configurations consistently occur within a narrow range of learning rates, whereas multiple combinations of the remaining architectural hyperparameters produce comparable predictive performance.**

**Fig. S4**. Slice plots illustrating the relationship between individual hyperparameters and the validation AUROC obtained during Bayesian hyperparameter optimization. Each point represents a single optimization trial and is colored according to the trial order. The learning rate exhibits the strongest association with predictive performance, whereas the remaining hyperparameters demonstrate comparatively low sensitivity within the explored search space.

**Fig. S5**. Pairwise contour plots illustrating the interactions among the optimized hyperparameters during Bayesian optimization. The contour color represents the validation AUROC (objective value), with darker regions indicating higher predictive performance. Each subplot depicts the relationship between a pair of hyperparameters explored during the optimization process, providing a comprehensive visualization of pairwise interaction effects across the hyperparameter search space.

**Fig. S6.** Representative attention weight visualizations for fifteen experimentally validated host–virus protein interactions recovered by the Local view model during external validation. Each panel (a–o) corresponds to one representative interaction selected from the high-confidence predictions summarized in Table 7. For each interaction, the left plot shows the attention entropy of the target host and viral proteins, whereas the right plot presents the ten neighboring proteins receiving the highest attention weights during graph message passing. Lower attention entropy indicates that the model concentrates on a smaller number of highly informative neighboring proteins, while higher entropy reflects a broader distribution of attention across the local neighborhood. These visualizations provide an interpretable representation of how the Local view model identifies informative local network contexts for host–virus interaction prediction.

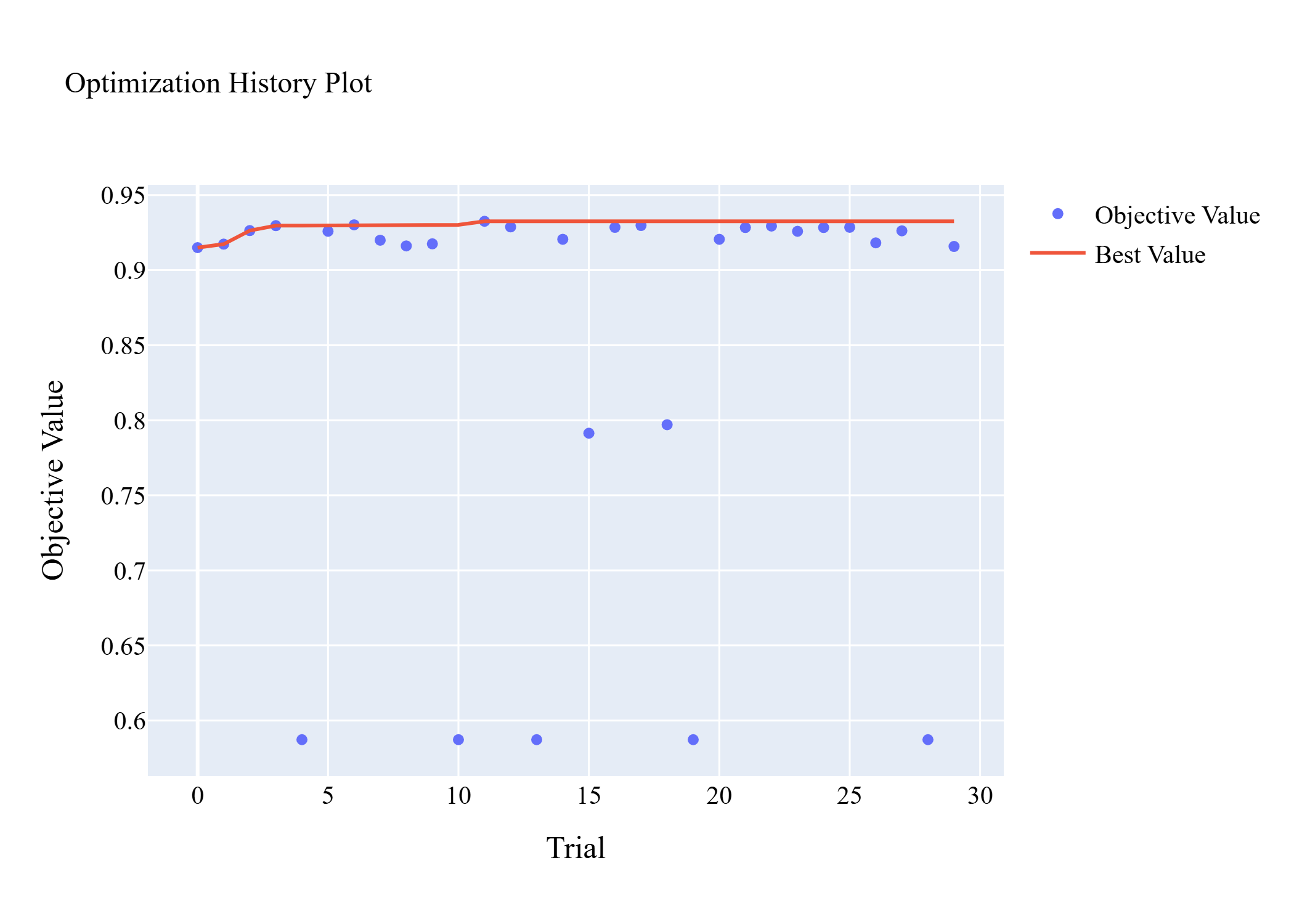

**Fig. S1**. Optimization history of Bayesian hyperparameter optimization using Optuna. Each blue point represents the validation AUROC obtained in an individual optimization trial, while the red line indicates the best objective value achieved up to each trial.

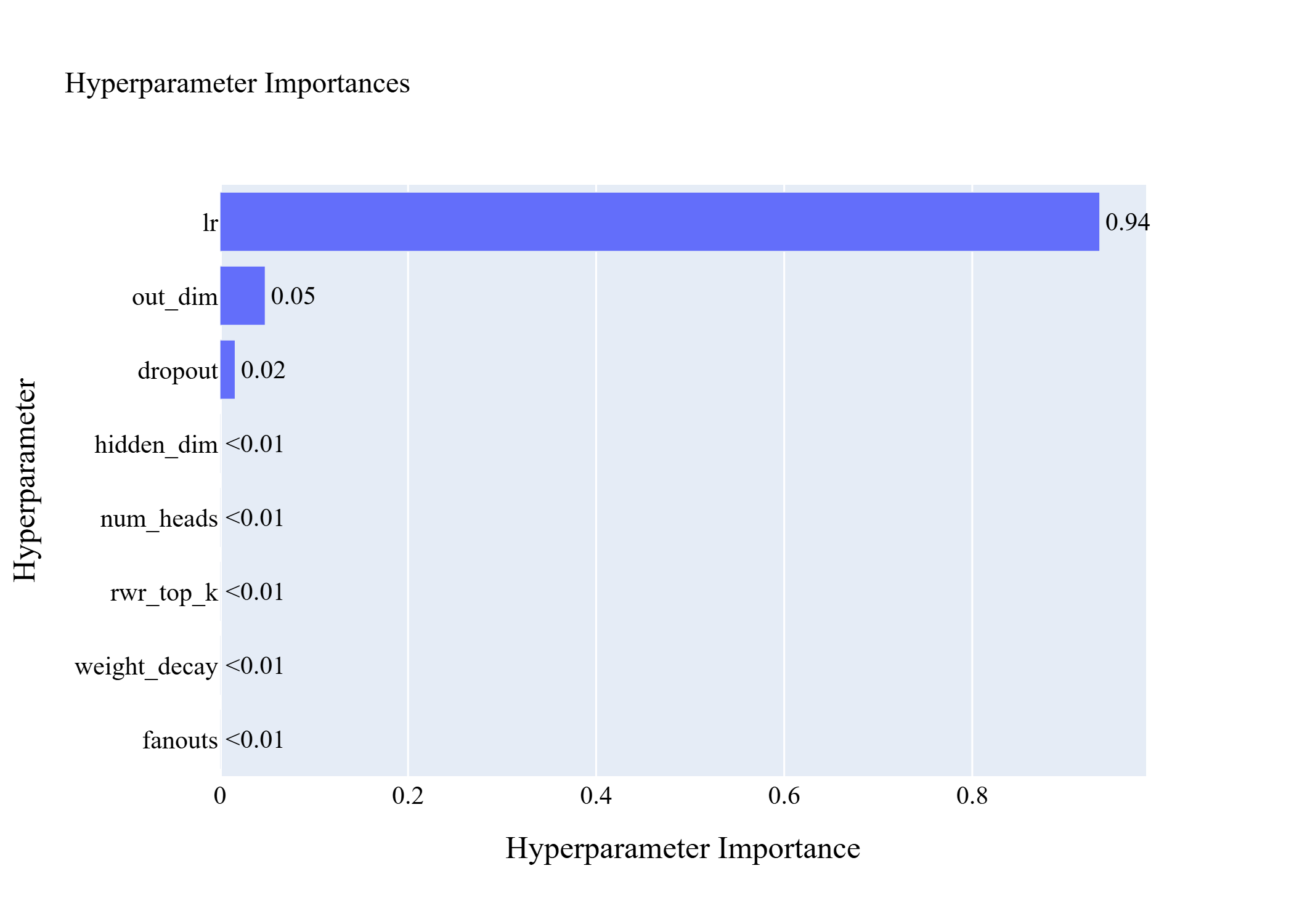

**Fig. S2**. Relative importance of optimized hyperparameters estimated by Optuna. Importance values indicate the contribution of each hyperparameter to the variation in validation AUROC across the 30 Bayesian optimization trials. The learning rate was the dominant factor influencing predictive performance, whereas the remaining hyperparameters exhibited comparatively low importance within the explored search space.

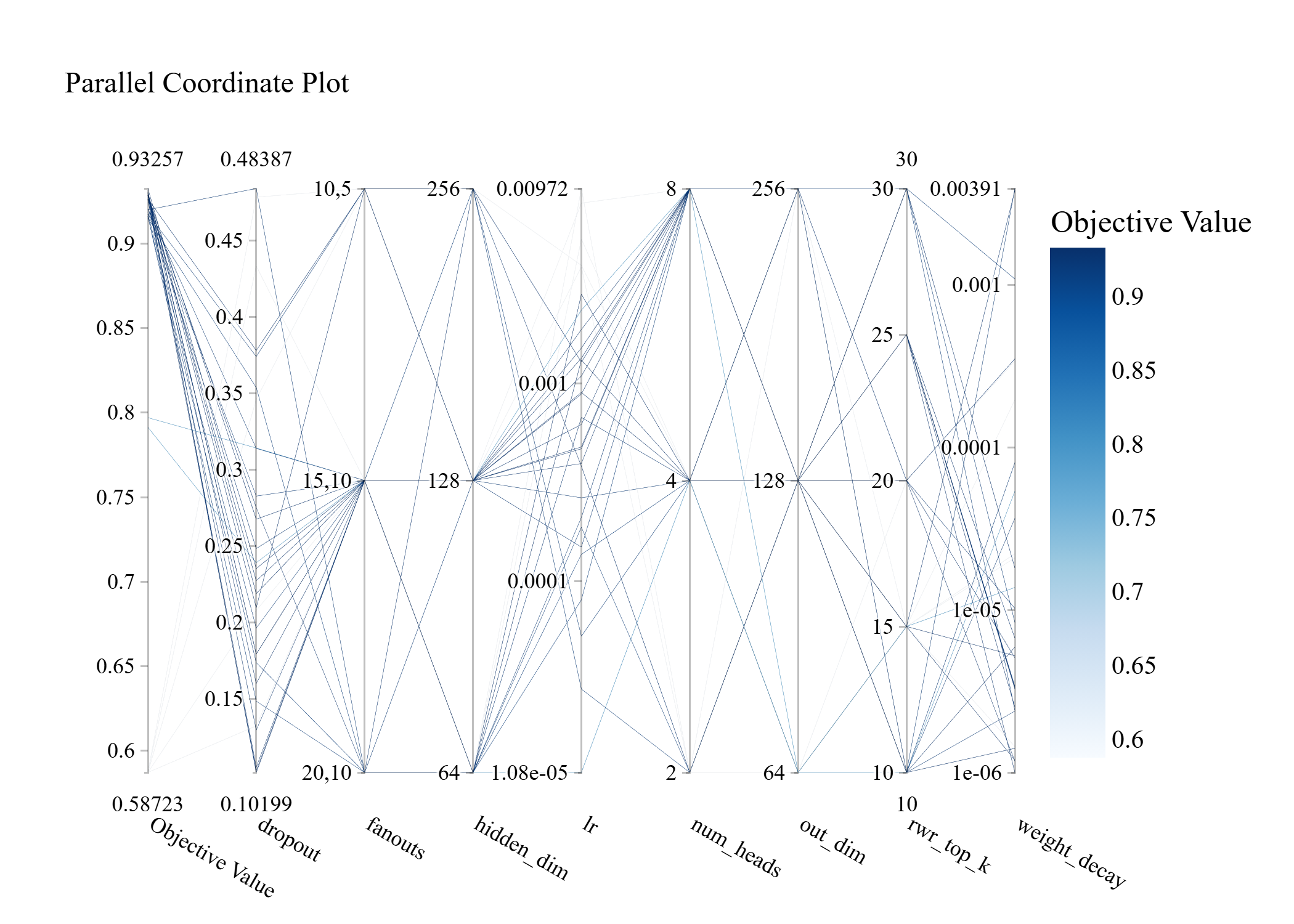

**Fig. S3. Parallel coordinate plot illustrating the hyperparameter configurations explored during Bayesian optimization. Each polyline represents one optimization trial, while the color indicates the validation AUROC (objective value), with darker lines corresponding to better-performing trials. The visualization shows that high-performing configurations consistently occur within a narrow range of learning rates, whereas multiple combinations of the remaining architectural hyperparameters produce comparable predictive performance.**

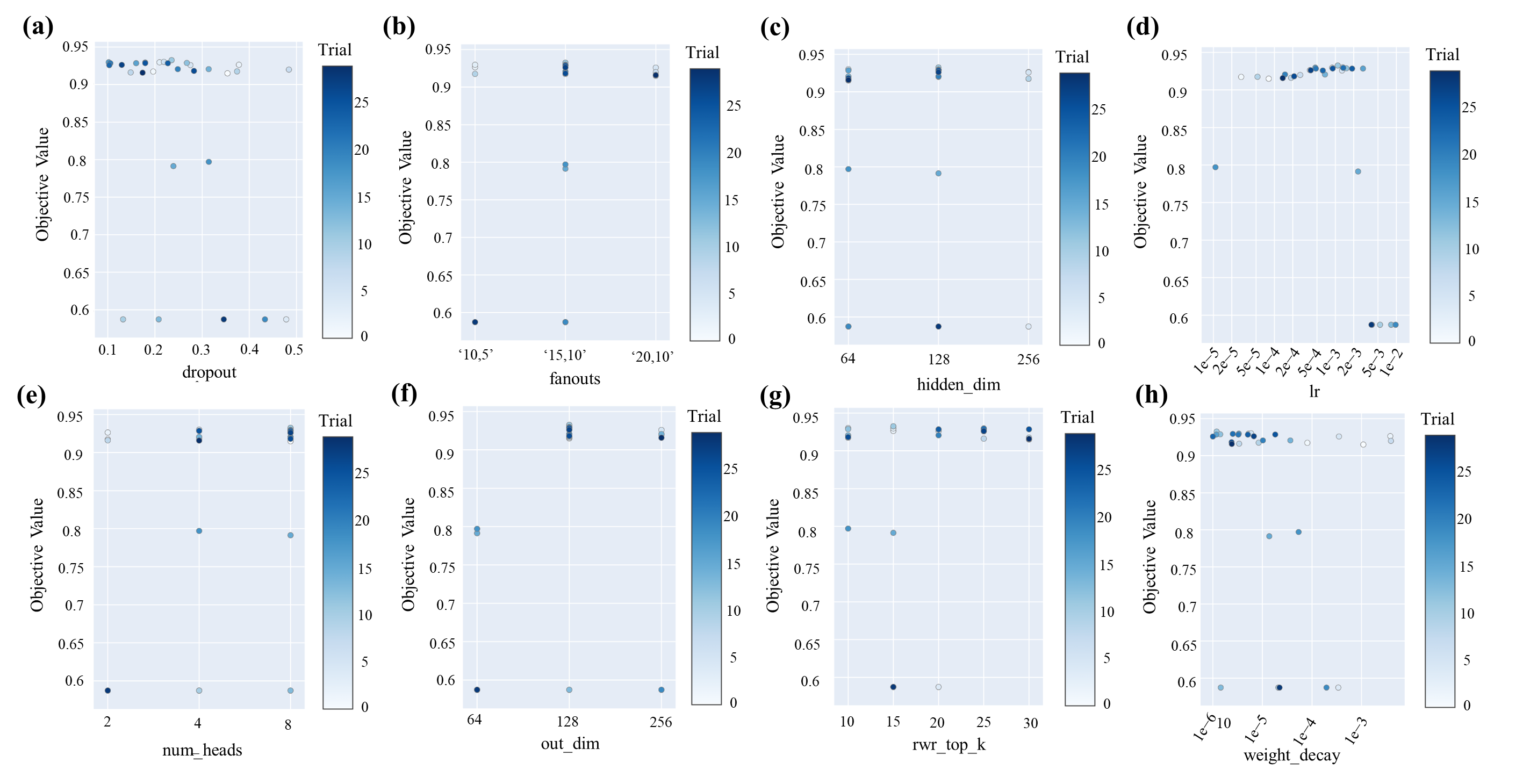

**Fig. S4**. Slice plots illustrating the relationship between individual hyperparameters and the validation AUROC obtained during Bayesian hyperparameter optimization. Each point represents a single optimization trial and is colored according to the trial order. The learning rate exhibits the strongest association with predictive performance, whereas the remaining hyperparameters demonstrate comparatively low sensitivity within the explored search space.

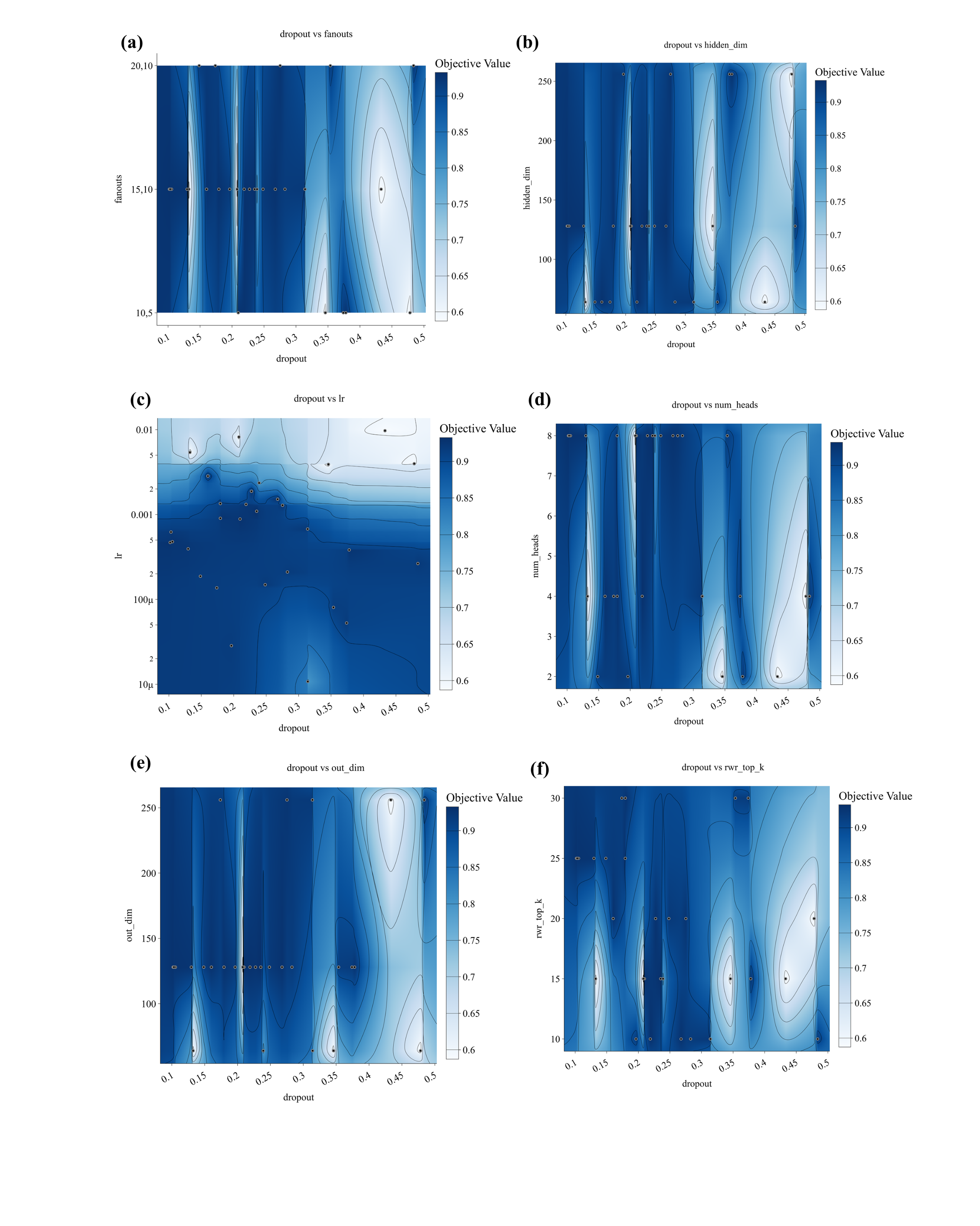

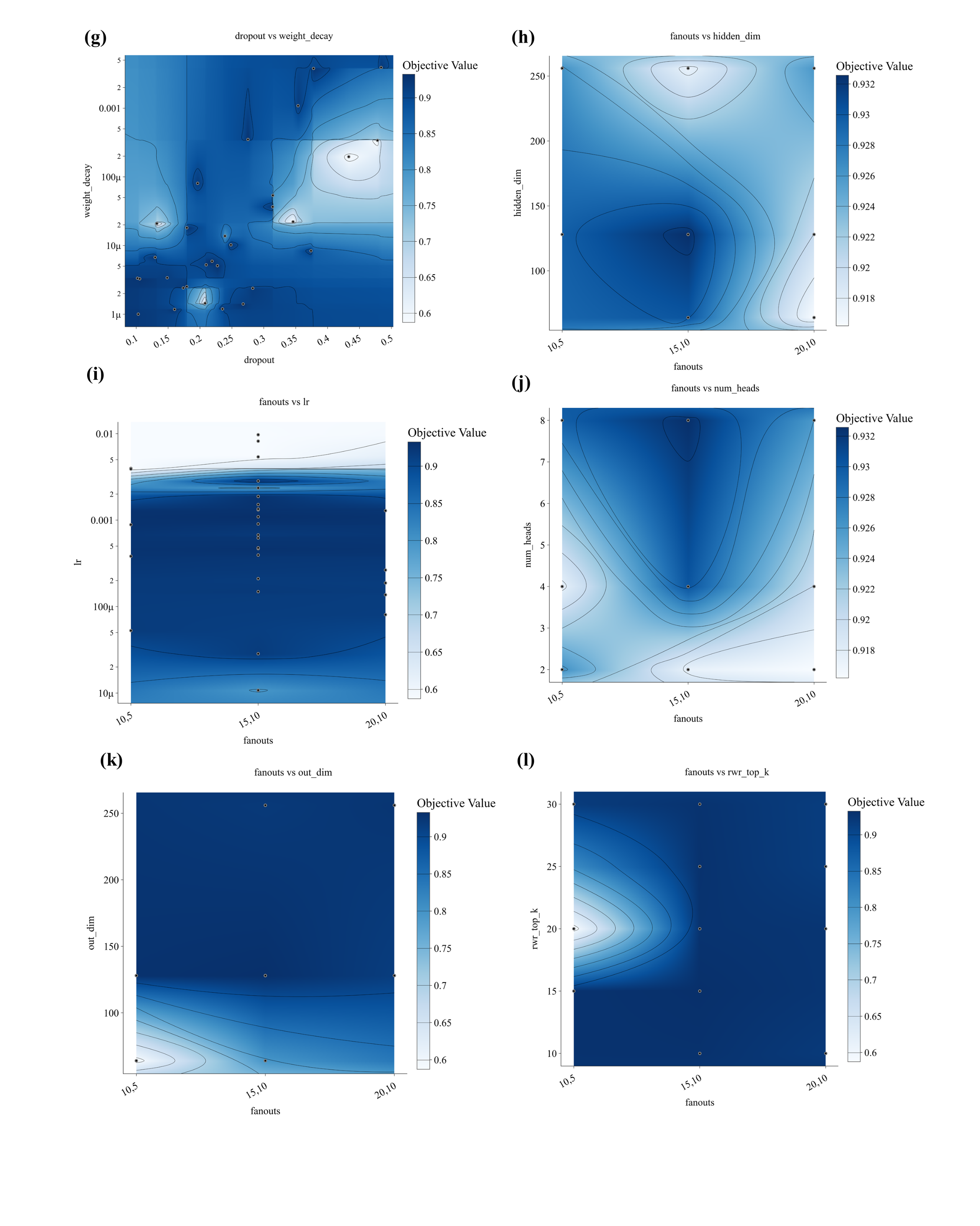

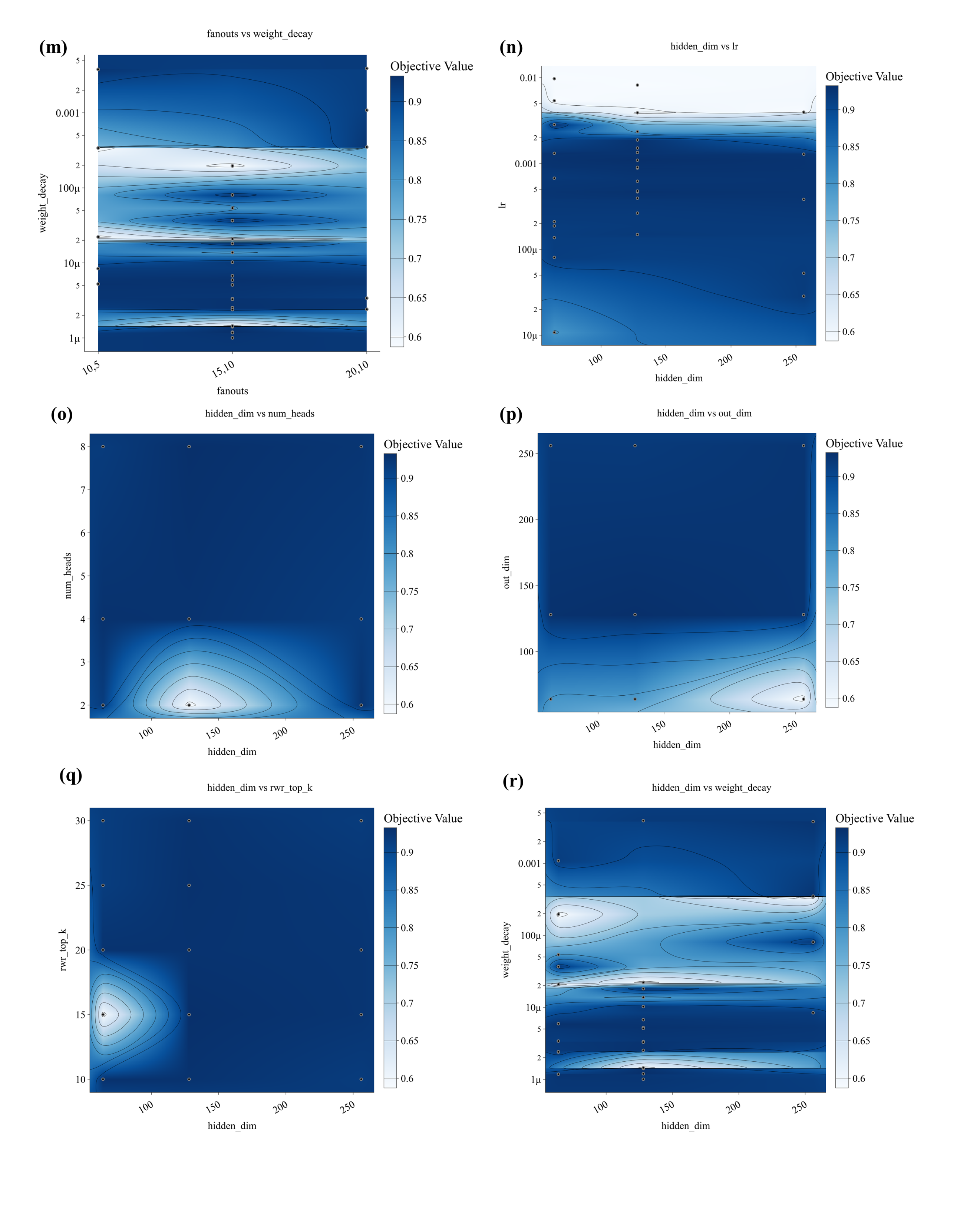

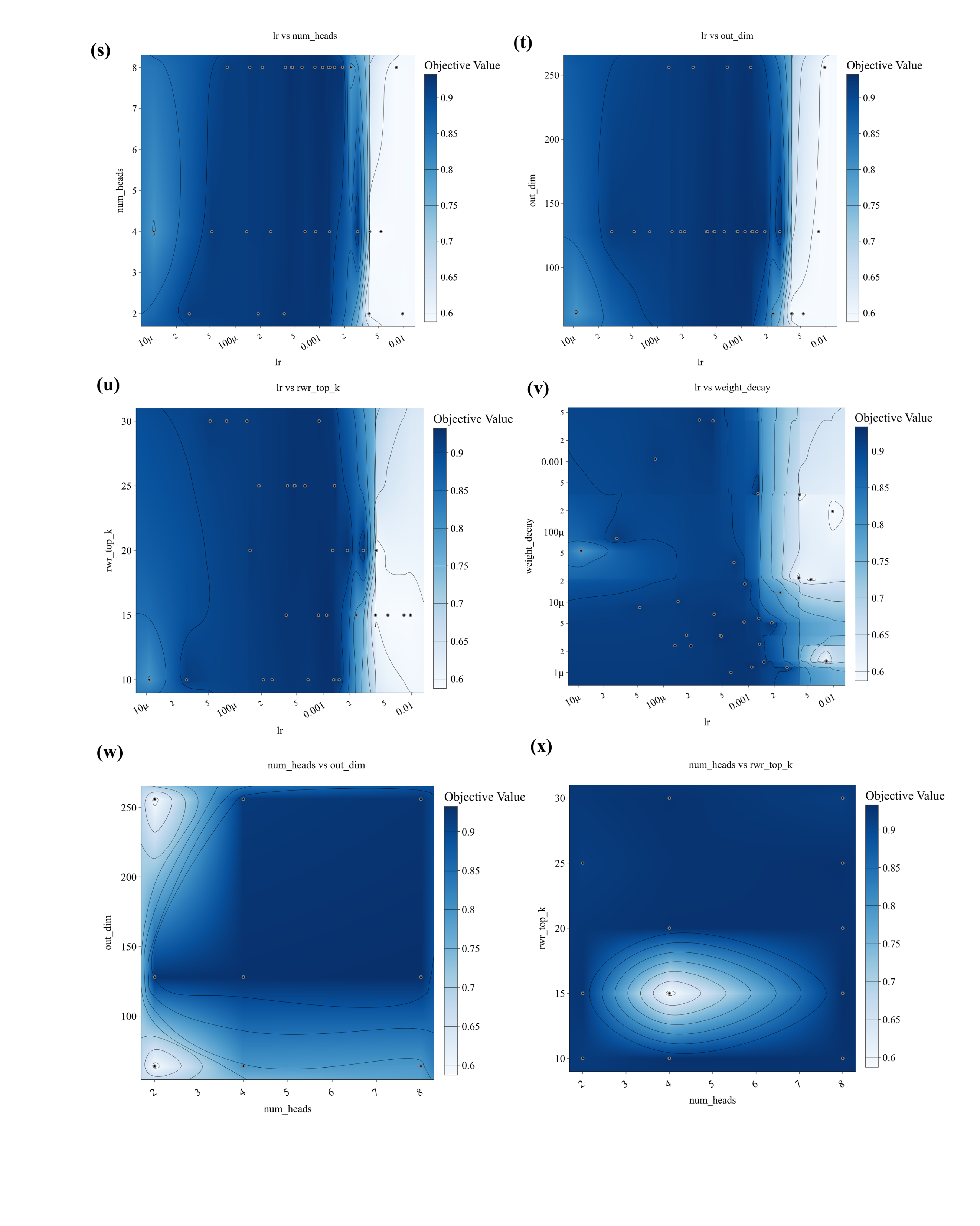

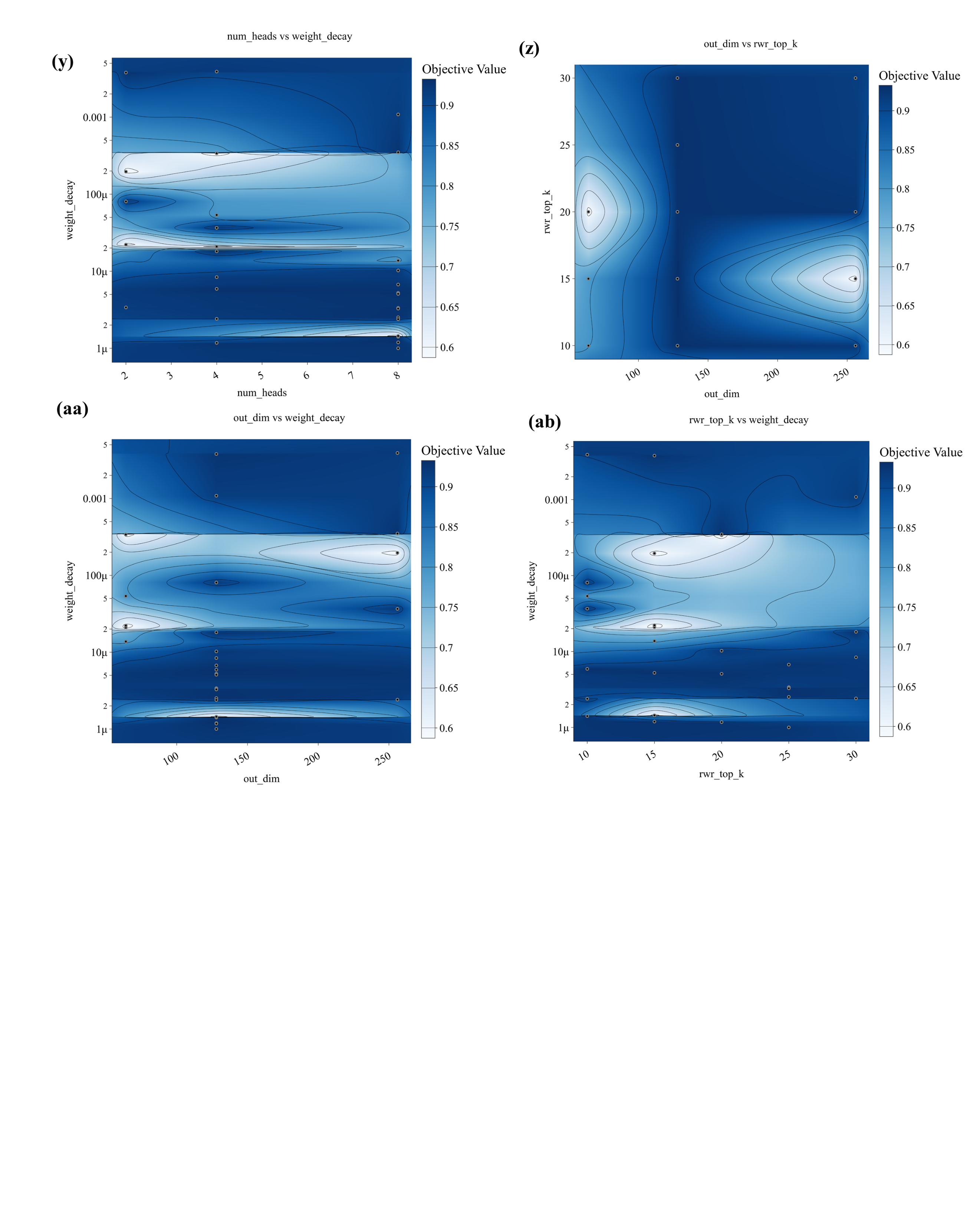

**Fig. S5**. Pairwise contour plots illustrating the interactions among the optimized hyperparameters during Bayesian optimization. The contour color represents the validation AUROC (objective value), with darker regions indicating higher predictive performance. Each subplot depicts the relationship between a pair of hyperparameters explored during the optimization process, providing a comprehensive visualization of pairwise interaction effects across the hyperparameter search space.

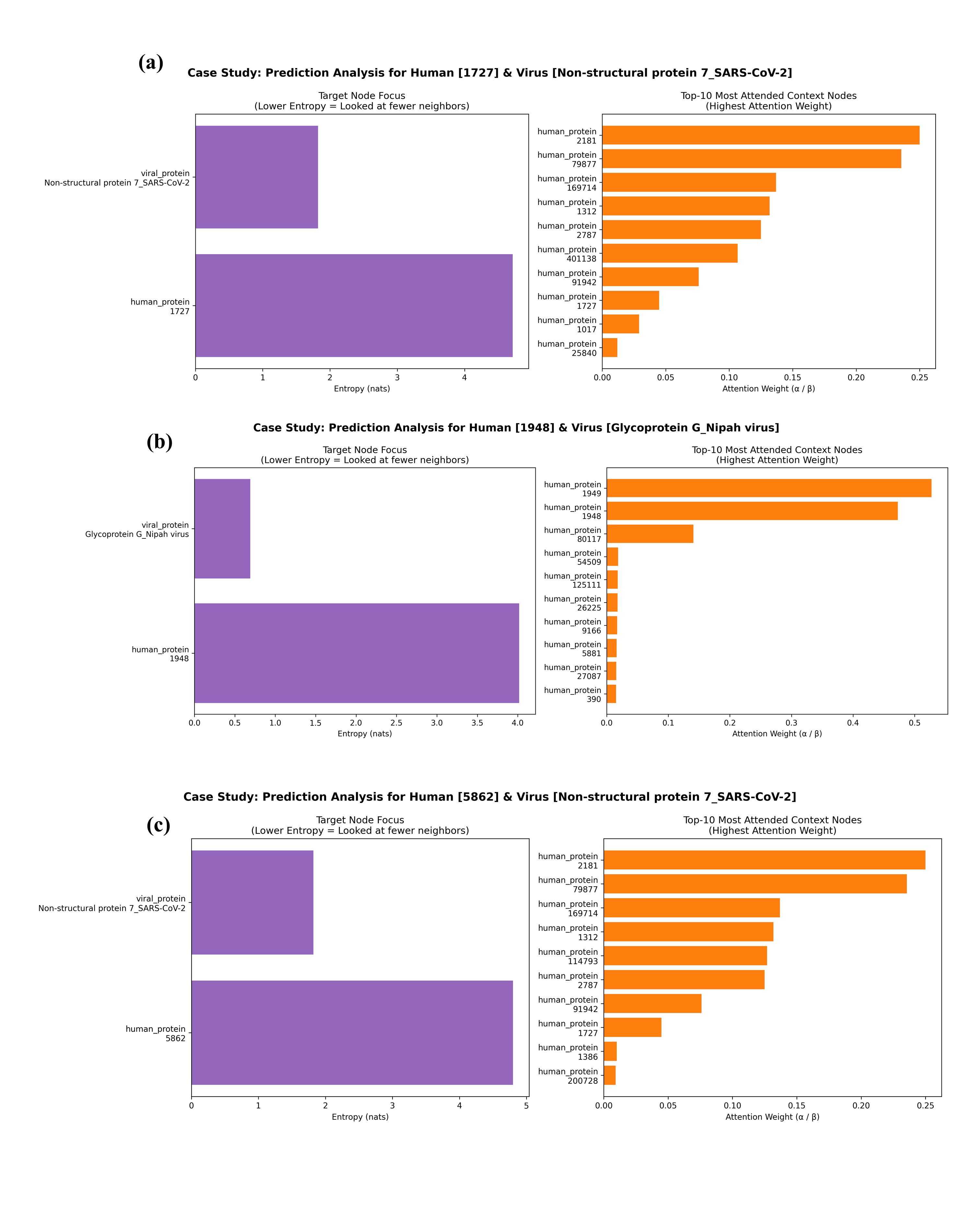

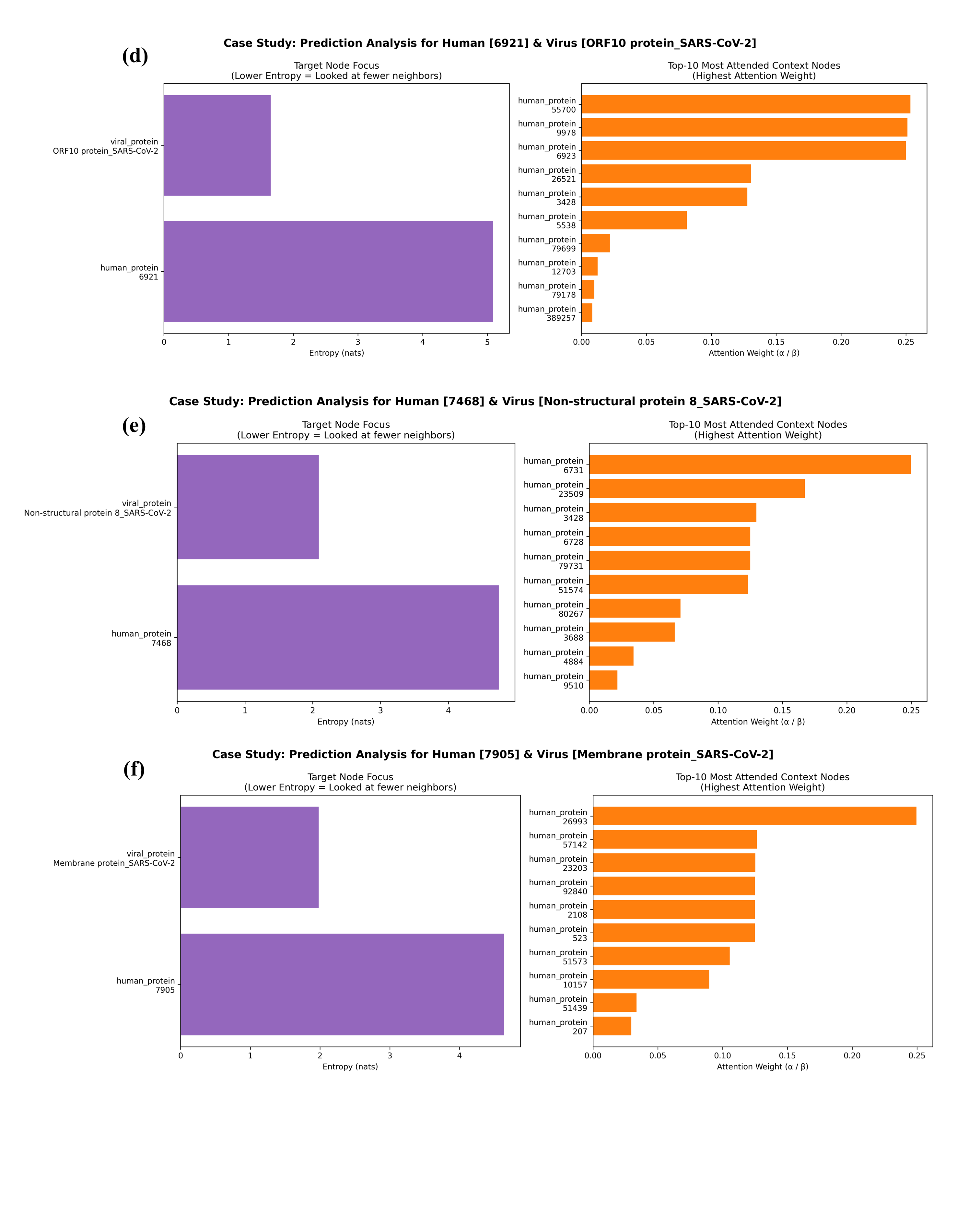

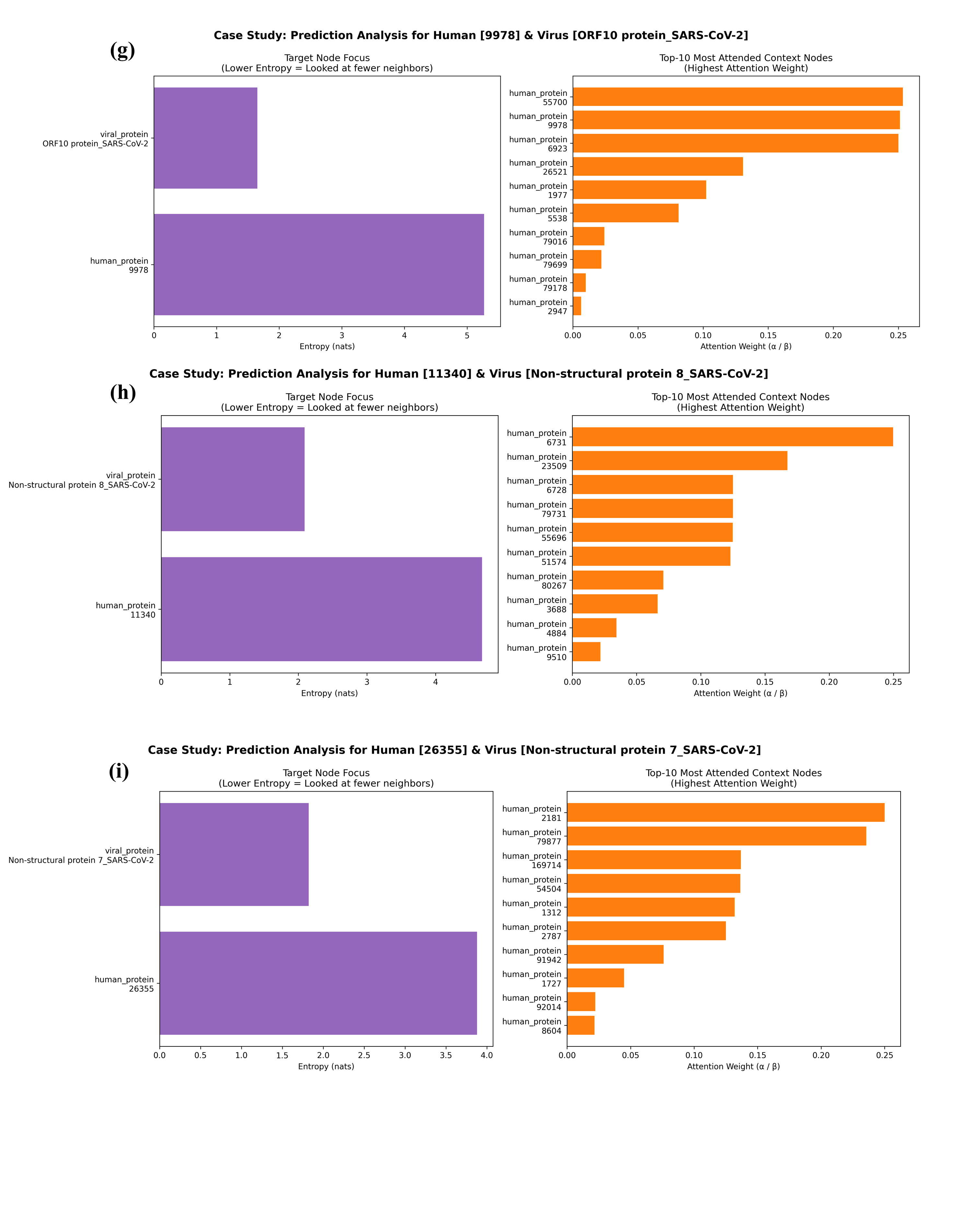

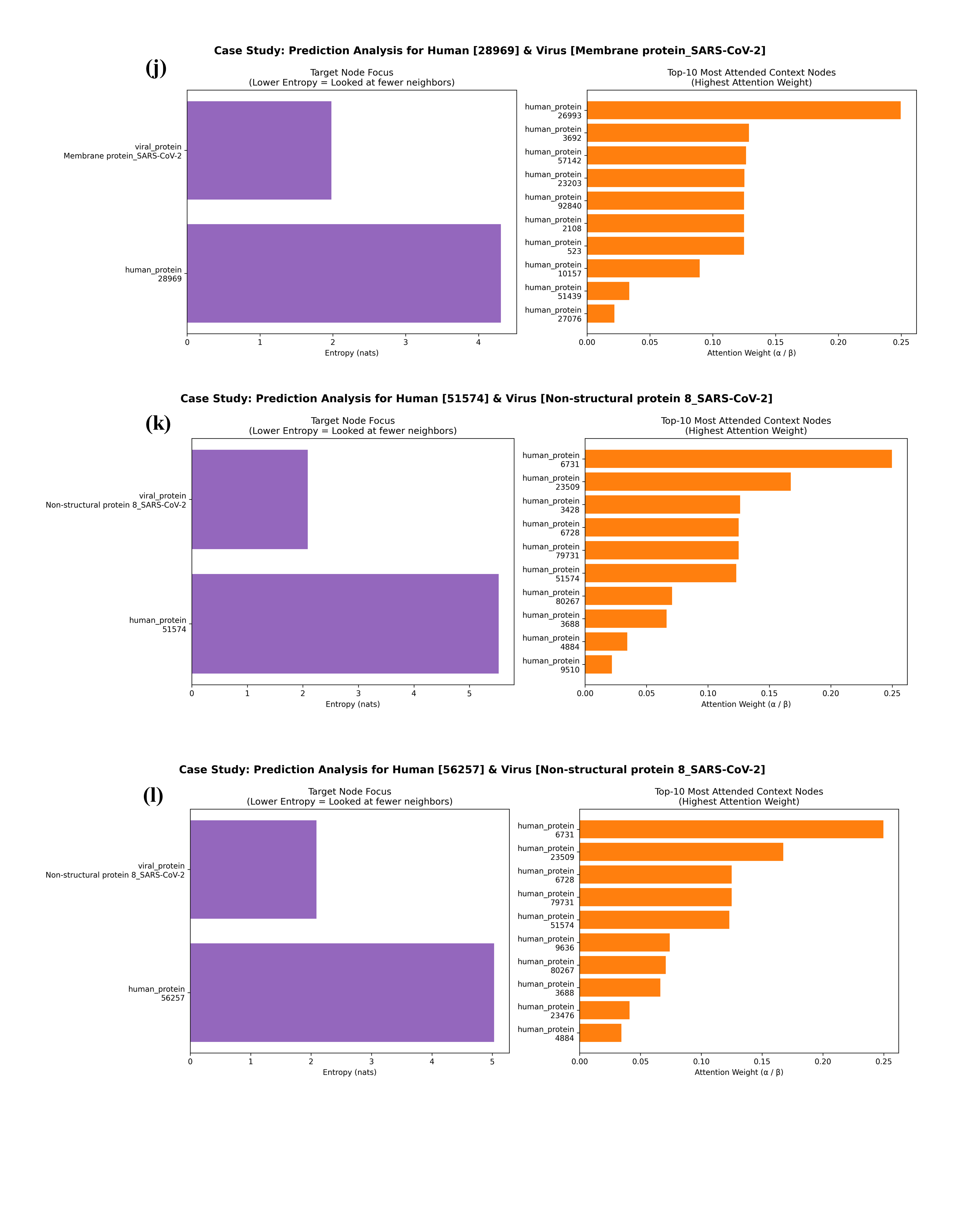

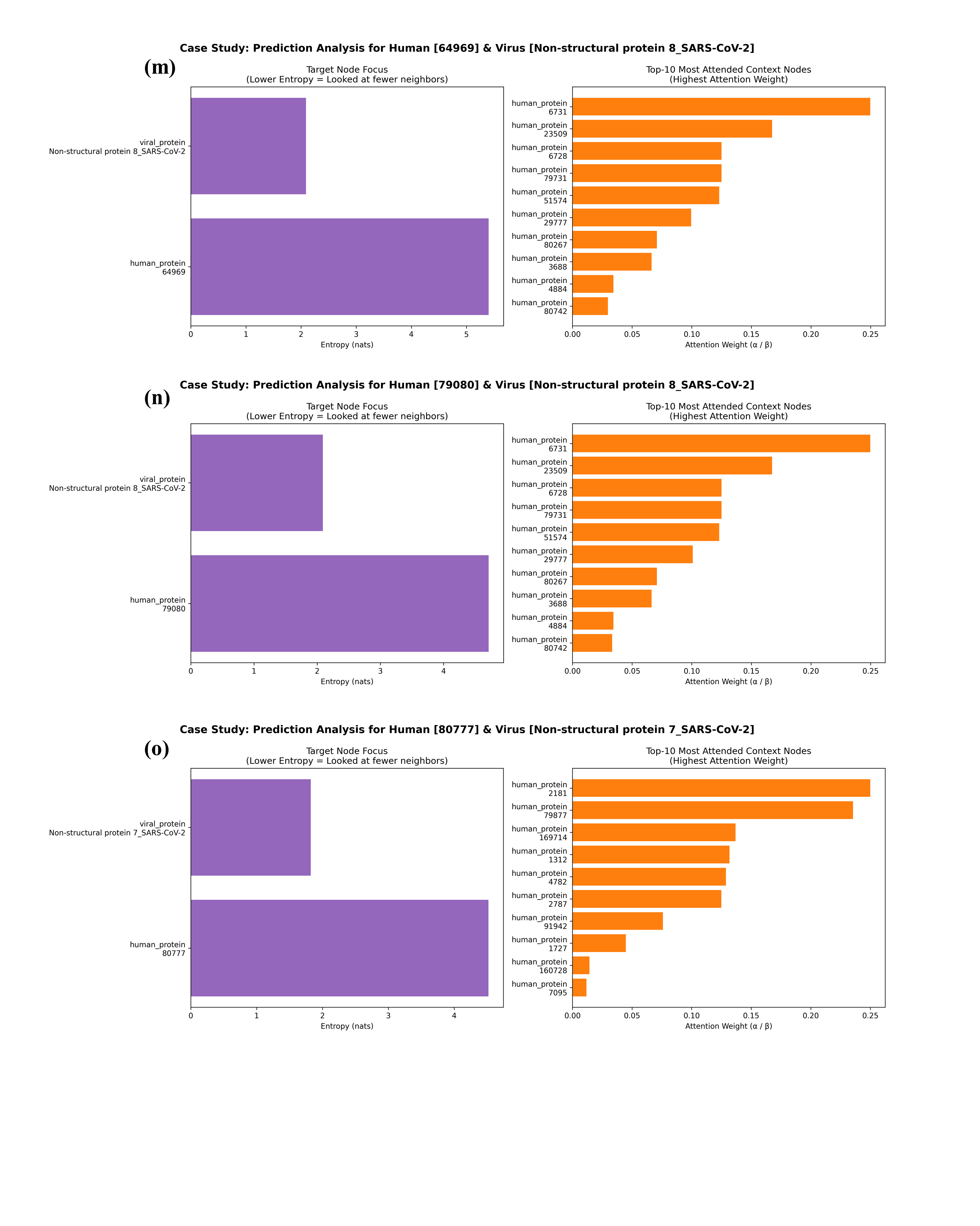

**Fig. S6.** Representative attention weight visualizations for fifteen experimentally validated host–virus protein interactions recovered by the Local view model during external validation. Each panel (a–o) corresponds to one representative interaction selected from the high-confidence predictions summarized in Table 7. For each interaction, the left plot shows the attention entropy of the target host and viral proteins, whereas the right plot presents the ten neighboring proteins receiving the highest attention weights during graph message passing. Lower attention entropy indicates that the model concentrates on a smaller number of highly informative neighboring proteins, while higher entropy reflects a broader distribution of attention across the local neighborhood. These visualizations provide an interpretable representation of how the Local view model identifies informative local network contexts for host–virus interaction prediction.
